# Epilepsy-related CDKL5 deficiency slows synaptic vesicle endocytosis in central nerve terminals

**DOI:** 10.1101/2022.03.15.484308

**Authors:** Christiana Kontaxi, Elizabeth C. Davenport, Peter C. Kind, Michael A. Cousin

## Abstract

Cyclin-dependent kinase-like 5 (CDKL5) deficiency disorder (CDD) is a severe early-onset epileptic encephalopathy resulting mainly from *de novo* mutations in the X-linked *CDKL5* gene. To determine whether loss of presynaptic CDKL5 function contributes to CDD, we examined synaptic vesicle (SV) recycling in primary hippocampal neurons generated from a *Cdkl5* knockout rat model. Using a genetically-encoded reporter, we revealed that CDKL5 is selectively required for efficient SV endocytosis. We showed that CDKL5 kinase activity is both necessary and sufficient for optimal SV endocytosis, since kinase-inactive mutations failed to correct endocytosis in *Cdkl5* knockout neurons, whereas the isolated CDKL5 kinase domain fully restored SV endocytosis kinetics. Finally, we demonstrated that CDKL5-mediated phosphorylation of amphiphysin 1, a putative presynaptic target, is not required for CDKL5-dependent control of SV endocytosis. Overall, our findings reveal a key presynaptic role for CDKL5 kinase activity and enhance our insight into how its dysfunction may culminate in CDD.

## Introduction

The majority of neuronal communication occurs at synapses, at which the presynapse contains an abundant number of synaptic vesicles (SVs) loaded with neurotransmitters that are generally released in response to neuronal activity. Following SV fusion, synchronized mechanisms of SV regeneration from the presynaptic plasma membrane guarantee the availability of readily releasable SVs upon repetitive firing and, hence, the fidelity of neurotransmission (Cousin, 2017; Soykan et al., 2016). Neurodevelopmental disorders affect more than 3 % of children worldwide and involve the disturbance of programmed brain development leading to cognitive, social and motor deficits with epileptic seizures being a frequently observed comorbidity (Parenti et al., 2020; Thapar et al., 2017). Mutations in several genes encoding for SV proteins have been identified as causal in the human condition (Baker et al., 2018; Dhindsa et al., 2015; Fassio et al., 2018; Salpietro et al., 2019; Serajee and Huq, 2015). In addition, multiple animal models that exhibit SV trafficking deficits display abnormalities reminiscent of neurodevelopmental conditions (Boumil et al., 2010; Di Paolo et al., 2002; Koch et al., 2011). Therefore, presynaptic dysfunction is emerging as a high-risk factor during neural development.

Cyclin-dependent kinase-like 5 (CDKL5) deficiency disorder (CDD) is a neurodevelopmental and epileptic encephalopathy that is primarily caused by *de novo* single-nucleotide mutations in the X-linked *CDKL5* gene (Fehr et al., 2013). CDD patients largely experience early-onset epileptic seizures and severe neurodevelopmental delay, in addition to a broad spectrum of other clinical manifestations. The human neuron-specific isoform of CDKL5 is a widely expressed serine/threonine kinase, consisting of an N-terminal catalytic domain followed by a long unstructured C-terminal tail (Kilstrup-Nielsen et al., 2012). CDKL5 has been implicated in various neuronal activities, including axon elongation (Nawaz et al., 2016), and synaptogenesis (Zhu et al., 2013). Furthermore, it is proposed to have synaptic roles, with hyperexcitability reported in both excitatory and inhibitory *Cdkl5* conditional knockout (KO) neurons (Tang et al., 2019; Tang et al., 2017). Likewise, upon loss of CDKL5, decreased spontaneous glutamate and GABA efflux is observed in cerebellar synaptosomes (Sivilia et al.,2016). However, a direct role for CDKL5 in SV recycling has not been explored.

Almost all pathogenic mutations in the *CDKL5* gene cluster within the region encoding its kinase domain (Hector et al., 2017), suggesting loss of its enzyme function may be key in CDD. Recently, a limited number of endogenous CDKL5 substrates were identified (Baltussen et al.,2018; Munoz et al., 2018), in addition to a series of *in vitro* targets (Baltussen et al., 2018; Sekiguchi et al., 2013). To date, the only *in vitro* presynaptic target of CDKL5 is amphiphysin 1 (Amph1), on the site serine 293 (S293) within a proline-rich domain (PRD). Amph1 is a cytosolic protein highly enriched in nerve terminals, where it acts as a hub during SV recycling via its multiple interaction domains, including its PRD (Wigge and McMahon, 1998; Wu et al.,2009). Importantly, S293 is a major *in vivo* phosphorylation site on Amph1 and is dephosphorylated during neuronal activity, indicating that it may be of high biological importance (Craft et al., 2008).

In the present study, we use a novel *Cdkl5* KO rat model (Simões de Oliveira et al. 2022) to examine SV recycling in CDKL5-deficient hippocampal neurons. Using the genetically-encoded fluorescent reporter synaptophysin-pHluorin (sypHy), we reveal that SV endocytosis is slower upon loss of CDKL5, but SV exocytosis remains unaffected. Following a molecular replacement strategy we demonstrate that the kinase activity of CDKL5 is both necessary and sufficient to correct dysfunction in SV endocytosis. Finally, we determined that the phosphorylation status of Amph1-S293 remains unaltered in CDKL5-null neurons, revealing that CDKL5 exerts its effect on SV endocytosis via a distinct presynaptic substrate. Taken together, our work reveals that CDKL5-mediated phosphorylation is critical for SV endocytosis efficiency, and that CDKL5 deficiency is responsible for presynaptic malfunction.

## Results

### Endogenous CDKL5 is sorted into the presynaptic terminal

CDKL5 is a ubiquitous neuronal protein kinase (Rusconi et al., 2011; Schroeder et al., 2019) however, its localisation at the nerve terminal has not been extensively addressed. To verify that CDKL5 is present in presynaptic terminals, and therefore in the correct location to influence SV recycling, a classical subcellular fractionation was performed. During this protocol, an adult rat brain was subjected to homogenisation and differential centrifugation to generate distinct subcellular fractions, including a crude synaptosome-(P2, mainly representing the presynapse with attached postsynaptic density) and an SV-enriched (LP2) fraction. Western blotting with a CDKL5-specific antibody (**Figure S1**) revealed that CDKL5 was present in the P2 fraction and enriched in the LP2 fraction, where the SV protein synaptophysin 1 (Syp1) also accumulated (**Figure 1A**). The relative absence of the postsynaptic marker, postsynaptic density 95 (PSD95), suggested that contamination of the LP2 fraction with postsynaptic elements was limited. Therefore, CDKL5 is present at presynaptic terminals and may associate with SVs, consistent with previous studies showing that CDKL5 colocalises with the presynaptic vesicular glutamate transporter 1 (VGLUT1) in mouse neurons (Ricciardi et al., 2012; Wang et al., 2021).

**Figure 1.**
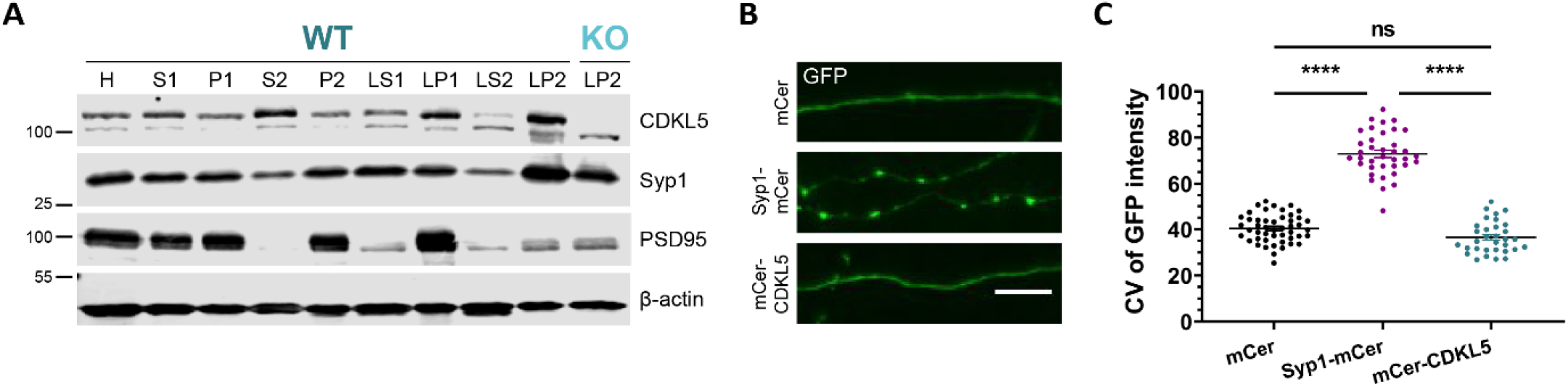
CDKL5 is present at nerve terminals. *(A)* Subcellular fractionation of adult rat brain for the crude purification of a synaptosomal (P2) and an SV (LP2) fraction and fractions representing other subcellular compartments (H, homogenate; P1, tissue debris, nuclei, and large myelin fragments; S2, microsomes, mitochondria, and synaptosomes; LP1, synaptic membrane, mitochondria, and myelin fragments; LS2, synaptosomal cytoplasm). An LP2 fraction from an adult CDKL5 KO rat brain was also generated. CDKL5 is enriched in the LP2 fraction (top band; 110 KDa). Synaptophysin 1 (Syp1) and postsynaptic density 95 (PSD95) were used as pre- and postsynaptic markers, respectively, and β-actin as loading control. (*B*) Mouse hippocampal neurons were transfected with either mCer, Syp1-mCer or mCer-CDKL5 at 8-9 DIV, fixed at 15 DIV, and stained for GFP. Examples of axonal segments of > 15 μm that were selected for coefficient of variation (CV) analysis are displayed. Scale bar, 5 μm. (*C*) The distribution pattern of CDKL5 was assessed by CV analysis of GFP fluorescence intensity along multiple > 15 μm axonal segments per field of view. Scatter plots indicate mean ± SEM. ns, not significant, \*\*\*\**p* < 0.0001 by one-way ANOVA followed by Tukey’s multiple comparison test. mCer *n* = 48, Syp1-mCer *n* = 37, mCer-CDKL5 *n* = 32 fields of view from 4 independent preparations of neuronal cultures.

To assess whether CDKL5 is targeted exclusively to nerve terminals or displays a more diffuse axonal distribution, we performed coefficient of variance (CV) analysis. Hippocampal neurons were transfected with either CDKL5 fused to the fluorescent protein mCerulean (mCer-CDKL5), Syp1-mCer or the empty mCer vector and were then immunolabelled for the presence of the fluorescent tag (**Figure 1B**). SV proteins, such as Syp1, are anticipated to have a punctate distribution along the axon and therefore a higher CV value. In contrast, lower CV values indicate a homogeneous distribution of a protein along the axon. In agreement, mCer-Syp1 displayed a localised distribution along the axon and a high CV value, in agreement with previous results (Gordon and Cousin, 2013). Quantification of the distribution profile of CDKL5 in axonal segments indicated a CV value similar to the empty mCer vector (**Figure 1C**). Therefore, CDKL5 is diffusely distributed along the axon, including presynaptic terminals.

### Loss of CDKL5 does not influence the levels of presynaptic proteins or the number of presynaptic boutons

We next investigated whether the absence of CDKL5 causes any defects in presynaptic stability since disruption of synapse stability/synaptogenesis may result in altered neuronal development. This was important to address, since dysregulation of protein levels in addition to altered synapse number have been reported in mice lacking CDKL5 (Della Sala et al., 2016; Ren et al., 2019; Schroeder et al., 2019; Tang et al., 2019). First, we examined whether expression of key presynaptic proteins was altered in rat CDKL5 KO neurons via Western blotting. Initially, we confirmed the absence of CDKL5 in lysates of KO neurons (**Figure 2A**). We then analysed a range of presynaptic molecules including proteins important for SV recycling, such as clathrin heavy chain (CHC), dynamin 1 (Dyn1), endophilin A1, and syndapin 1; integral SV proteins, such as Syp1, VGLUT1, and the v-type proton ATPase subunit B (ATP6V1B2); and phosphoproteins that have been implicated in the regulation of SV endocytosis, such as the protein kinases glycogen synthase kinase 3 (GSK3) and Akt (Clayton et al., 2010; Ferreira et al., 2021; Smillie and Cousin, 2012). These latter enzymes were of particular interest, since the PI3K/GSK3/Akt pathway has been one of the most perturbed signalling cascades in CDKL5 deficiency model systems (Amendola et al., 2014; Jiang et al.,2019; Wang et al., 2012). This analysis revealed that the absence of CDKL5 did not significantly alter the total protein level of any candidate, or the phosphorylation status (and thus activity) of either GSK3 or Akt when compared to wild-type (WT) controls (**Figure 2A**). Therefore CDKL5 KO neurons do not display overt alterations in presynaptic proteins or signalling cascades.

**Figure 2.**
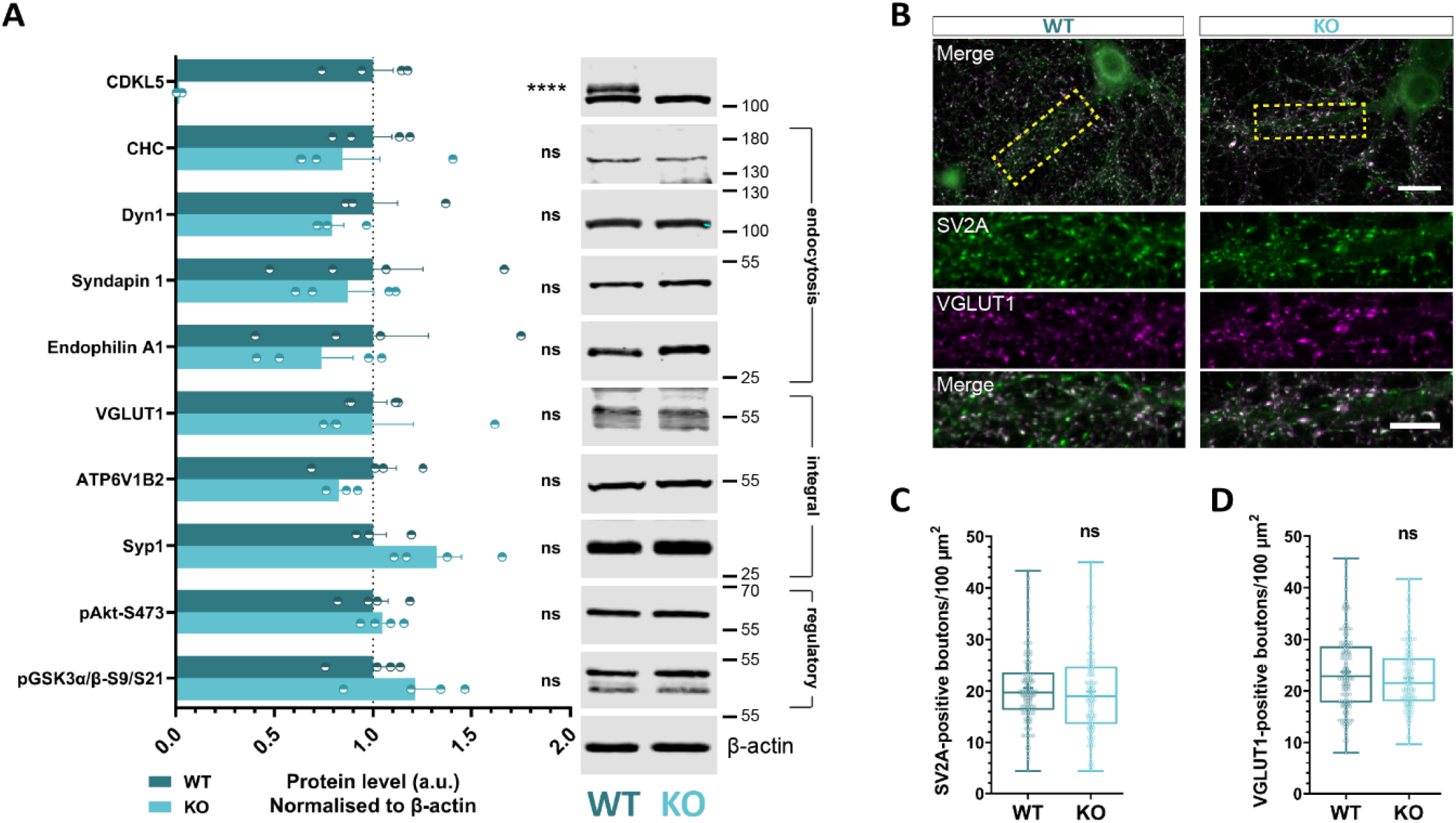
Loss of CDKL5 does not alter presynaptic protein levels or the number of presynaptic boutons. *(A)* Hippocampal neuronal lysates at 14-15 DIV were analysed for different presynaptic proteins, including SV endocytosis proteins, integral SV proteins, and phosphoproteins regulating SV endocytosis. Quantification of the total band intensity normalised to β-actin revealed no difference for any of these proteins in the absence of CDKL5. Bars indicate mean ± SEM. ns, not significant, \*\*\*\**p* < 0.0001 by unpaired two-tailed t test. WT *n* = 4, KO *n* = 4 neuronal lysates from 4 independent preparations of neuronal cultures. (*B*) Hippocampal neuronal cultures derived from WT and CDKL5 KO rats were fixed at 15 DIV and stained for the presynaptic proteins SV2A and VGLUT1. The number of positive puncta was counted in (50 × 15) μm^2^ selections along processes (dashed yellow boxes) for both markers. Scale bar, 20 μm (neurons), 10 μm (processes). (*C*) Quantification of SV2A-positive boutons and (*D*) VGLUT1-positive boutons per 100 μm^2^ along WT and CDKL5 KO dendrites. Box plots present median with IQR indicating min to max whiskers. ns, not significant by Mann Whitney two-tailed t test, + indicates mean value. WT *n* = 144, KO *n* = 142 neurons from 3 independent preparations of neuronal cultures.

Next, we investigated whether the lack of CDKL5 led to a reduced number of presynaptic terminals. To achieve this, WT and CDKL5 KO neurons were double-stained for two distinct presynaptic markers, synaptic vesicle protein 2A (SV2A) and VGLUT1, to assess the number of presynaptic boutons and excitatory presynaptic subtypes, respectively. There were no genotype-specific differences in SV2A- and VGLUT1-positive puncta along neuronal processes (**Figure 2B**). Therefore, there is no effect of the absence of CDKL5 on either the number of total or excitatory presynaptic terminals (**Figure 2C, D**). Overall, this data reveals that the formation and maintenance of nerve terminals in rat primary neuronal cultures is not affected upon CDKL5 deficiency.

### Loss of CDKL5 impairs SV regeneration but does not influence SV exocytosis

The presynaptic localisation of CDKL5 suggests that CDKL5 is implicated in SV recycling. Indeed, phenotypes reported in mice lacking CDKL5, such as altered frequency of spontaneous and miniature postsynaptic currents (mPSCs) (Tang et al., 2017; Wang et al.,2021), and aberrant paired-pulse facilitation (Tang et al., 2019), indicate that CDKL5 deficiency results in defects in synaptic transmission that could be due to dysfunctional SV recycling. To determine this, we used the genetically-encoded reporter sypHy, in which a pH-sensitive form of GFP, ecliptic pHluorin (pKa ~7.1), is inserted into an intravesicular loop of Syp1 (Granseth et al., 2006; Miesenbock et al., 1998). The fluorescence of sypHy is dictated by the pH of its immediate environment, with fluorescence being quenched in the acidic SV lumen, unquenched upon stimulus-dependent SV exocytosis and exposure to the cell surface, and re-quenched following endocytosis and SV acidification (**Figure 3A**). To determine the potential contribution of CDKL5 to SV recycling across a range of stimulus intensities, primary hippocampal neurons derived from CDKL5 KO rats or WT littermate controls were transfected with sypHy and stimulated with action potential (AP) trains of either 5 Hz or 10 Hz (both 300 APs) or 40 Hz (400 APs) (**Figure 3B, E, H**). To quantify for the extent of activity-dependent SV exocytosis, the amount of sypHy fluorescence during stimulation was measured as a proportion of the total fluorescence within the presynapse revealed by perfusion with NH4Cl that allows for an estimation of the total recycling SV pool. We found that the extent of SV exocytosis remained unaltered between genotypes across all stimulation frequencies investigated (**Figure 3C, F, I**). To confirm this phenotype, we next measured the rate of sypHy fluorescence increase during prolonged stimulation (10 Hz for 90 s) in the presence of bafilomycin A1. Bafilomycin A1 is a V-type ATPase inhibitor, and therefore removes any potential contribution from SV endocytosis to the sypHy response during the stimulation by blocking SV acidification (Sankaranarayanan and Ryan, 2001). When this experiment was performed, no difference was observed in either the rate of the sypHy fluorescence increase (SV exocytosis rate) or the extent of the sypHy response (SV recycling pool size) between WT and CDKL5 KO neurons (**Figure S2A, B, C**). Therefore, SV exocytosis is not altered upon CDKL5 loss.

**Figure 3.**
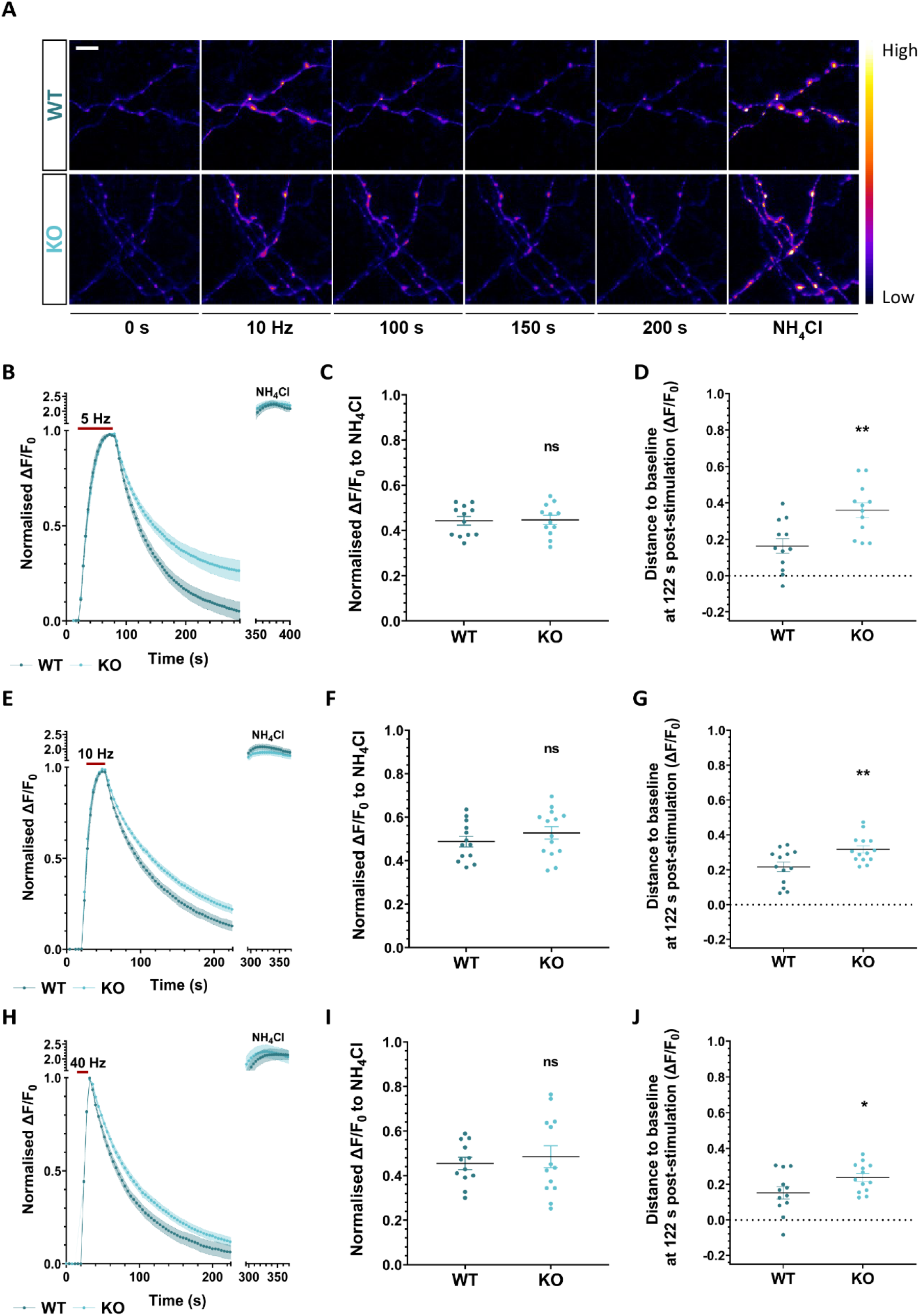
Loss of CDKL5 impairs the kinetics of SV endocytosis but not SV exocytosis. Primary hippocampal neurons from WT and CDKL5 KO rats were transfected with sypHy at 8-9 DIV and used at 13-14 DIV. *(A)* Example responses from sypHy-expressing axons that were subjected to 300 APs at 10 Hz and perfused with NH4Cl solution 3 min after termination of stimulation. Representative image slices were selected from the time-course that was recorded from both WT (top) and CDKL5 KO neuronal cultures (bottom). Scale bar, 5 μm. SypHy response from neurons stimulated with either 300 APs at 5 Hz (*B*), 300 APs at 10 Hz (*E*), or 400 APs at 40 Hz (*H*) (red bars) normalised to the stimulation peak (ΔF/F0). (*C*, *F, I*) SypHy fluorescence (ΔF/F0) at stimulation peak when total sypHy response was normalised to NH_4_Cl. (*D*, *G, J*) sypHy fluorescence (ΔF/F_0_) measuring the distance from baseline at 122 s post-stimulation. Scatter plots indicate mean ± SEM. ns, not significant, \**p* < 0.05, \*\**p* < 0.01 by unpaired two-tailed *t* test. (*B*-*D*) WT *n* = 12, KO *n* = 12 coverslips from 4 independent preparations of neuronal cultures. (*E*-*G*) WT *n* = 13, KO *n* = 14 coverslips from 4 independent preparations of neuronal cultures. (*H*-*J*) WT *n* = 12, KO *n* = 13 coverslips from 4 independent preparations of neuronal cultures.

We next focused on SV endocytosis, in which protein kinases perform an important role (Clayton et al., 2010; Tan et al., 2003). As acidification is a rapid process when compared to rate-limiting SV endocytosis (Atluri and Ryan, 2006; Egashira et al., 2015; Granseth et al.,2006), monitoring the sypHy fluorescence decay after stimulation can be used to estimate SV endocytosis kinetics (Sankaranarayanan and Ryan, 2000). To quantify the kinetics of SV retrieval, the sypHy stimulation peak was normalised, and the amount of sypHy remaining to be retrieved 2 minutes after termination of stimulation was measured. This parameter was used for consistency across protocols, since in specific cases the decay kinetics were not mono-exponential (rendering time constant measurements redundant). CDKL5 KO neurons consistently displayed slower SV endocytosis across all frequencies examined when compared to WT, suggesting that CDKL5 is important for optimal SV endocytosis (**Figure 3D, G, J**). Interestingly, the requirement for CDKL5 appeared to be more prominent at lower stimulation frequencies.

To confirm that this phenotype was due to slowed SV endocytosis and not dysfunctional SV acidification, we determined the kinetics of SV acidification using an acid-pulse protocol (Granseth et al., 2006). In this protocol, an impermeant acid buffer (pH 5.5) is perfused immediately after stimulation to quench all surface sypHy, which exclusively reveals the sypHy signal inside recently retrieved SVs (where the quenching rate can be calculated). In this protocol WT and CDKL5 KO neurons expressing sypHy are perfused with acid buffer both prior to stimulation (to reveal an initial baseline) and immediately after stimulation (10 Hz, 30 s, to reveal the quenching rate inside SVs) (**Figure S2D**). No significant difference in the SV acidification rate in neurons lacking CDKL5 compared to WT neurons was apparent (**Figure S2E**), confirming that the slowing in the post-stimulus sypHy fluorescence decay in CDKL5 KO neurons was due to impaired SV endocytosis.

CDD is a disorder of early life, and a therefore key question to address is whether defects can be rescued by the re-introduction of the gene, or whether the altered circuit activity in its absence renders gene correction redundant. To address this in our system, we determined whether expression of WT CDKL5 in KO neurons could correct SV endocytosis deficits. Both CDKL5 KO and WT littermate controls were co-transfected with sypHy and either mCer-CDKL5 or an empty mCer vector and stimulated with either 300 APs at 10 Hz or 400 APs at 40 Hz. Analysis of the post-stimulus sypHy response showed that expression of mCer-CDKL5 fully restored the kinetics of SV endocytosis after 10 Hz stimulation and partially after 40 Hz. Importantly, mCer-CDKL5 overexpression had no impact on SV endocytosis kinetics in WT neurons, indicating that increased levels of the protein kinase had no dominant negative effect (**Figure 4A-D**). Thus, expression of CDKL5 can restore presynaptic defects observed in KO neurons.

**Figure 4.**
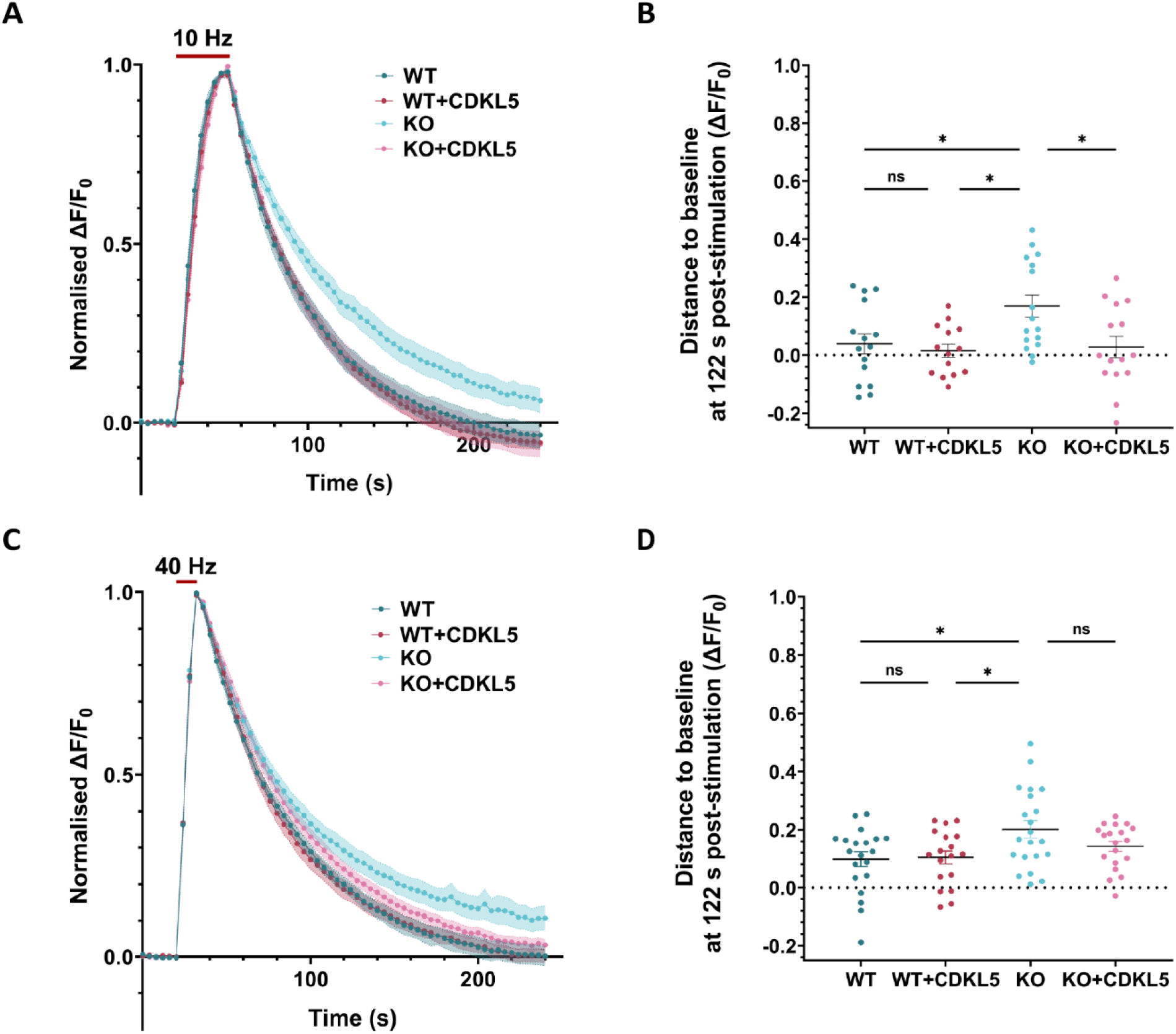
CDKL5 rescues the kinetics of SV endocytosis in CDKL5-deficient neurons. Primary hippocampal neurons from WT and CDKL5 KO rats were co-transfected with sypHy and mCer (WT, dark turquoise; KO, dark pink) or mCer-CDKL5 (WT+CDKL5, light turquoise; KO+CDKL5, light pink) at 8-9 DIV and used at 13-14 DIV. (*A*,*C*) sypHy response from neurons stimulated (red bar) with either 300 APs at 10 Hz (*A*) or 400 APs at 40 Hz (*C*) normalised to the stimulation peak and (*B*,*D*) sypHy fluorescence measuring the distance from baseline at 122 s post-stimulation. (*B*) Scatter plots indicate mean ± SEM. ns, not significant, *\*p* < 0.05 by two-way ANOVA followed by Tukey’s multiple comparison test. WT *n* = 15, WT+CDKL5 *n* = 14, KO *n* = 16, KO+CDKL5 *n* = 15 coverslips from 4 independent preparations of neuronal cultures. (*D*) Scatter plots indicate mean ± SEM. ns, not significant, \**p* < 0.05 by two-way ANOVA followed by Tukey’s multiple comparison test. WT *n* = 20, WT+CDKL5 *n* = 18, KO *n* = 21, KO+CDKL5 *n* = 19 coverslips from 5 independent preparations of neuronal cultures.

### CDD-related mutants of CDKL5 fail to rescue SV endocytosis impairment

As stated above, in CDD all identified pathogenic missense mutations are found within the kinase domain suggesting the disorder is due to loss of its enzymatic function (Hector et al.,2017; Munoz et al., 2018). To determine whether the protein kinase activity of CDKL5 is essential for its role in SV endocytosis, we investigated the ability of two mutant forms of full-length CDKL5 to restore function in CDKL5 KO neurons. The CDKL5 mutants were 1) K42R (a catalytically-inactive form of the enzyme that cannot bind ATP (Lin et al., 2005)), and 2) R178P, a mutation reported in CDD patients of both sexes with severe neurological features (Elia et al., 2008; Nemos et al., 2009) (**Figure 5A**). CDKL5 KO neurons were co-transfected with sypHy and either WT CDKL5 or one of the CDKL5 mutants and SV endocytosis kinetics were monitored following stimulation with either 300 APs at 10 Hz or 400 APs at 40 Hz (**Figure 5B, D**). WT CDKL5 fully restored SV endocytosis kinetics after both stimulation trains, as observed previously. In contrast, neither of the CDKL5 mutants were able to correct the SV endocytosis defect (**Figure 5C, E**). The absence of rescue was not due to their low expression, since this was equivalent to the exogenously-expressed WT enzyme (**Figure S3**). These data reveal that the protein kinase activity of CDKL5 is essential for optimal SV endocytosis kinetics and also associates CDKL5 pathology with defective SV recycling.

**Figure 5.**
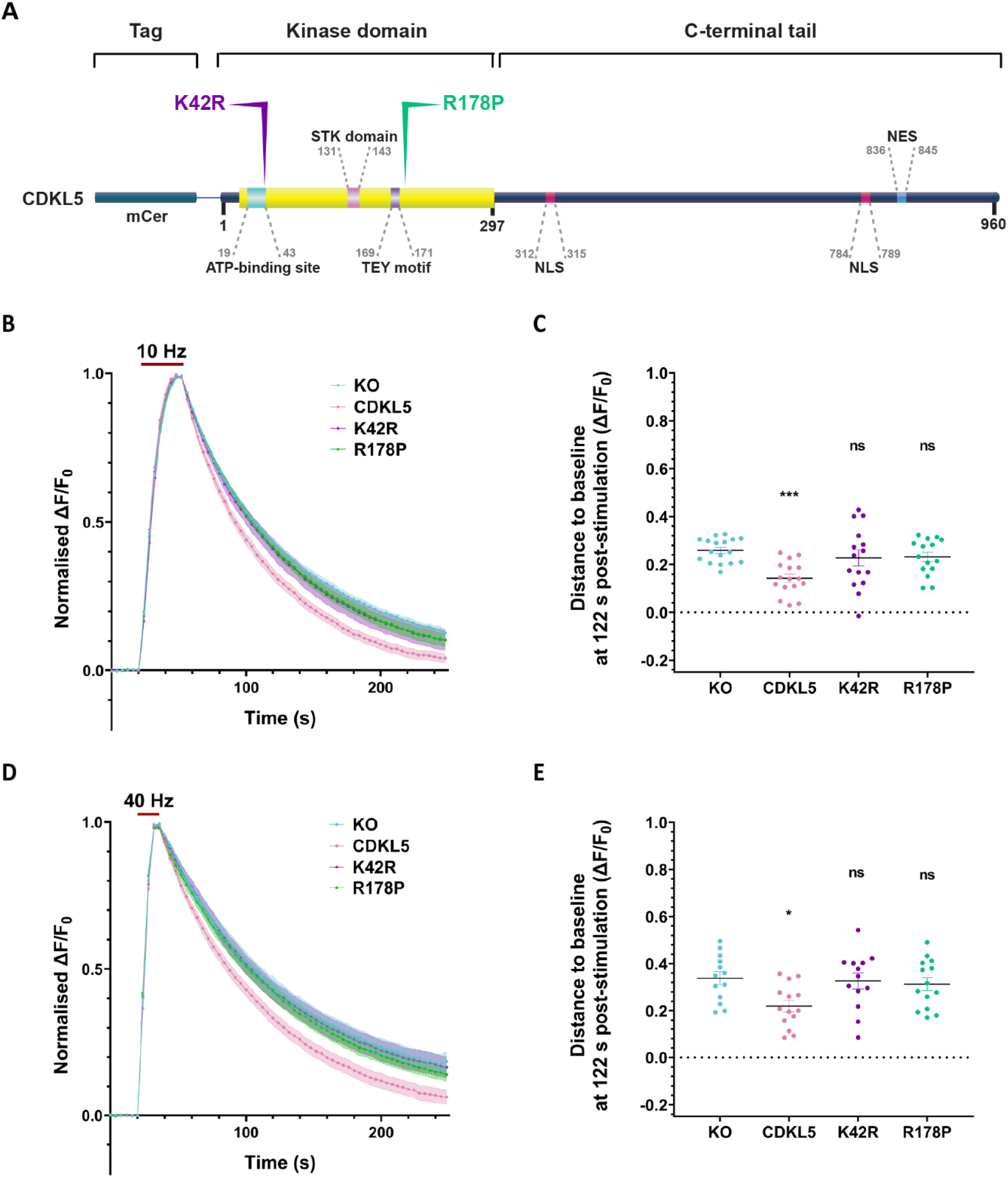
Point mutations within the CDKL5 kinase domain cannot correct SV endocytosis kinetics in CDKL5 KO neurons. (*A*) Schematic representation of the structural domains of CDKL5. Point mutations were introduced into the kinase domain including K42R, within the ATP-binding region, and R178P adjacent to the TEY motif. All constructs were tagged with mCer at their N-termini. (*B*-*E*) Primary hippocampal neurons from CDKL5 KO rats were co-transfected with sypHy and mCer (KO, light turquoise), mCer-CDKL5 (CDKL5, light pink), K42R (purple) or R178P (green) at 8-9 DIV and used at 13-14 DIV. (*B*,*D*) sypHy response from neurons stimulated (red bar) with either 300 APs at 10 Hz (*B*) or 400 APs at 40 Hz (*D*) normalised to the stimulation peak. (*C*,*E*) SypHy fluorescence measuring the distance from baseline at 122 s post-stimulation. (*C*) Scatter plots indicate mean ± SEM. ns, not significant, \*\*\**p* < 0.001 by one-way ANOVA followed by Dunnett’s multiple comparison test. KO *n* = 17, CDKL5 *n* = 16, K42R *n* = 15, R178P *n* = 15 coverslips from 5 independent preparations of neuronal cultures. (*E*) Scatter plots indicate mean ± SEM. ns, not significant, \**p* < 0.05 by one-way ANOVA followed by Dunnett’s multiple comparison test. KO *n* = 13, CDKL5 *n* = 14, K42R *n* = 13, R178P *n* = 14coverslips from 4 independent preparations of neuronal cultures.

### The kinase activity of CDKL5 is necessary and sufficient for optimal SV endocytosis

We have revealed an essential requirement for the enzymatic activity of CDKL5 in SV endocytosis. However a key question to address is whether this activity is both necessary and sufficient to correct SV endocytosis dysfunction in CDKL5 KO neurons. To address this, we examined whether expression of the isolated protein kinase domain was sufficient to correct presynaptic function in CDKL5 KO neurons. To determine this, we generated mCer-tagged deletion mutants of CDKL5 comprising either the kinase domain (ΔC; aa 1-297) or the C-terminal tail (Δ kinase; aa 298-960) (**Figure 6A**). Primary cultures of hippocampal CDKL5 KO neurons were co-transfected with sypHy and either full-length CDKL5 or one of the deletion mutants. Double immunostaining of primary cultured hippocampal neurons for GFP and endogenous CDKL5 suggested that Δkinase was expressed to higher levels than WT, whereas ΔC could not be quantified due to the absence of an antibody epitope (**Figure S3**). SV endocytosis kinetics were assessed by monitoring sypHy fluorescence after stimulation with 300 APs at 10 Hz or 400 APs at 40 Hz (**Figure 6B, D**). We observed that the isolated kinase domain was sufficient to rescue SV endocytosis kinetics similarly to full-length CDKL5 at both stimulus intensities (**Figure 6C, E**). In contrast, the isolated C-terminus could not, suggesting that this region cannot support SV endocytosis in the absence of the protein kinase domain. Therefore, the ability of the isolated CDKL5 protein kinase domain to correct presynaptic function reveals that it is both necessary and sufficient to rescue SV endocytosis, and that the C-terminal tail is dispensable for this role.

**Figure 6.**
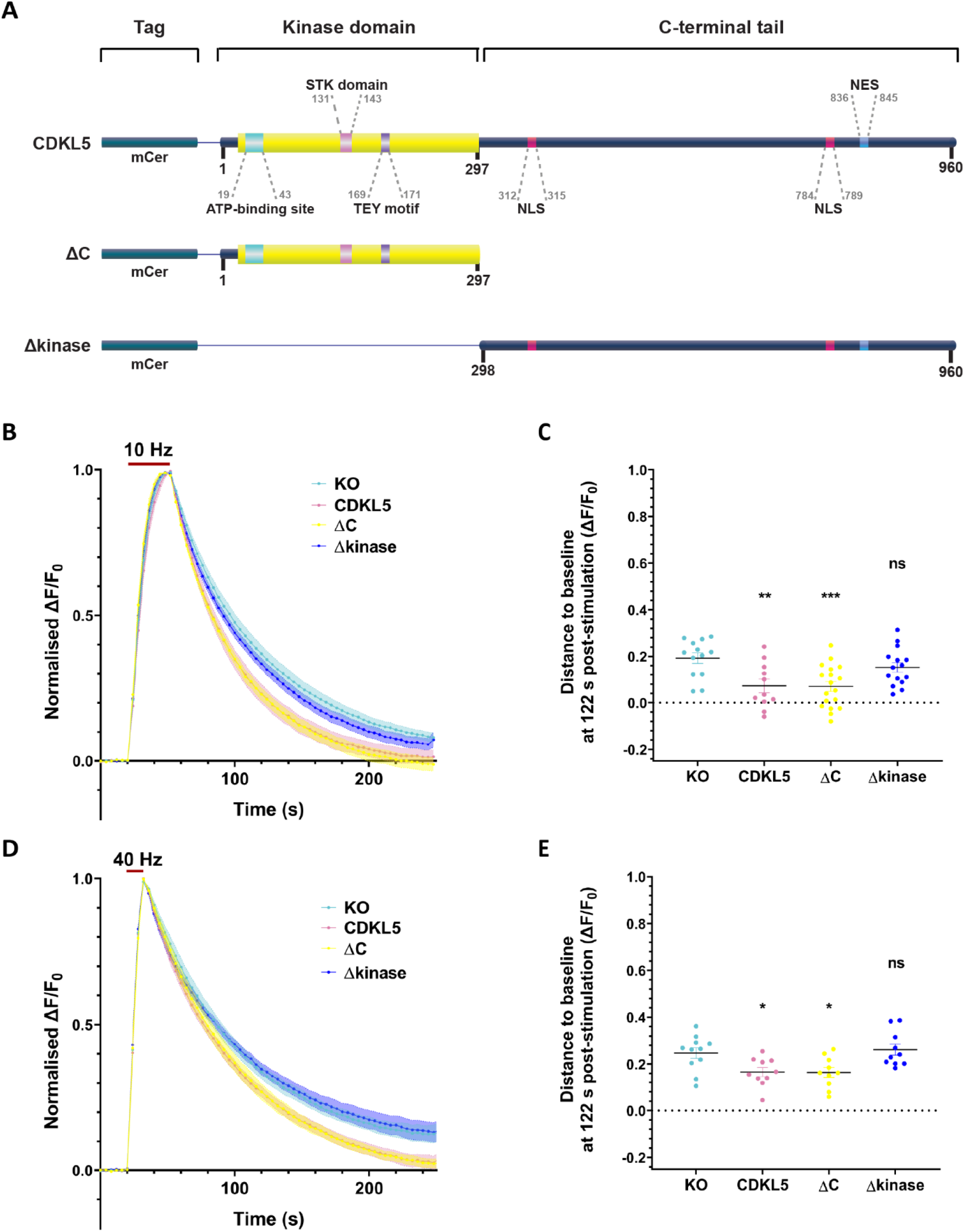
The CDKL5 kinase domain is sufficient to restore the SV endocytosis kinetics in CDKL5 KO neurons. (*A*) Schematic representation of the structural domains of CDKL5. Truncated versions of CDKL5 were generated comprising either the kinase domain (ΔC) or the C-terminal tail (Δkinase). All constructs were tagged with mCer at their N-termini. (*B*-*E*) Primary hippocampal neurons from CDKL5 KO rats were co-transfected with sypHy and mCer (KO, light turquoise), mCer-CDKL5 (CDKL5, light pink), the kinase domain (ΔC, yellow) or the C-terminal tail (Δkinase, blue) at 8-9 DIV and used at 13-14 DIV. (*B*,*D*) sypHy response from neurons challenged (red bar) with either 300 APs at 10 Hz (*B*) or 400 APs at 40 Hz (*D*) normalised to the stimulation peak. *(C,E)* sypHy fluorescence measuring the distance from baseline at 122 s post-stimulation. (*C*) Scatter plots indicate mean ± SEM. ns, not significant, \*\**p* < 0.01, \*\*\**p* < 0.001 by one-way ANOVA followed by Dunnett’s multiple comparison test. KO *n* = 13, CDKL5 *n* = 11, ΔC *n* = 18, Δkinase *n* = 15 coverslips from 4 independent preparations of neuronal cultures. (*E*) Scatter plots indicate mean ± SEM. ns, not significant \**p* < 0.05 by one-way ANOVA followed by Dunnett’s multiple comparison test. KO *n* = 11, CDKL5 *n* = 10, ΔC *n* = 10, Δkinase *n* = 10 coverslips from 3 independent preparations of neuronal cultures.

### CDKL5-mediated phosphorylation at Amph1-S293 is not required for SV regeneration

Since the kinase activity of CDKL5 is necessary for optimal SV endocytosis, this suggests that there is at least one CDKL5 substrate at the presynapse that mediates this role. The only candidate presynaptic target of CDKL5 that has been identified so far is Amph1, from *in vitro* studies (Katayama et al., 2015; Sekiguchi et al., 2013). To determine whether Amph1 may be a *bona fide* CDKL5 substrate, we first examined the ability of these two proteins to interact with each other, as it would be anticipated for an enzyme to interact with its substrates, even transiently. We demonstrated reciprocal co-immunoprecipitation of Amph1 and CDKL5 from rat brain lysates (**Figure 7A**). This indicates that CDKL5 binds to Amph1 *in vivo,* and hence supports that Amph1 may be a CDKL5 substrate.

**Figure 7.**
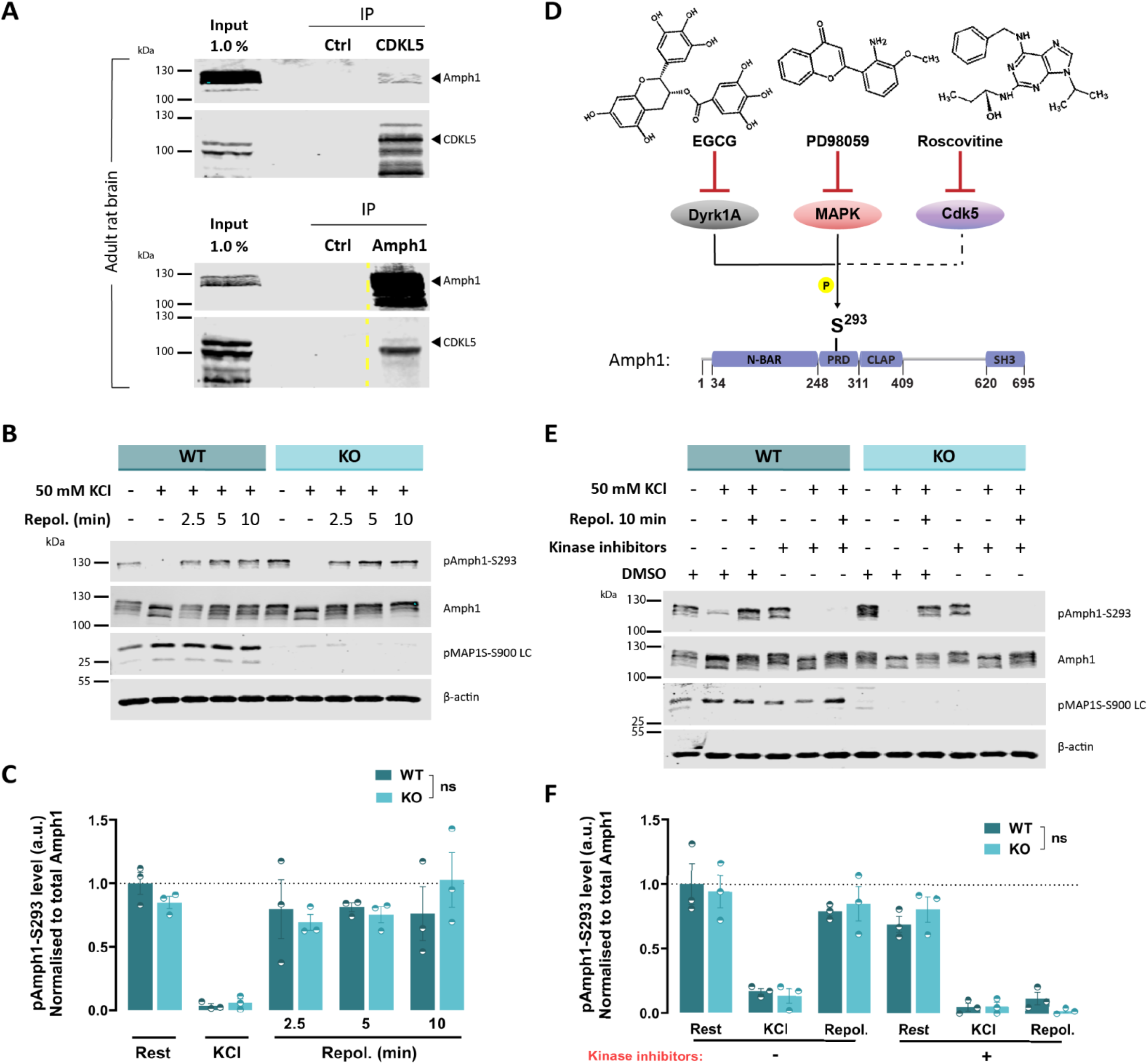
Amph1-S293 is phosphorylated independently of CDKL5. (*A*) Co-immunoprecipitation from adult rat brain lysates of Amph1 and CDKL5 with CDKL5 and Amph1 antibodies, respectively. Dashed lines indicate cropped images. (*B*) Primary hippocampal neurons at 14-15 DIV from WT and CDKL5 KO rats were depolarised with 50 mM KCl for 2 min and allowed to repolarise for 2.5, 5, and 10 min, respectively. Neurons were analysed for pAmph1-S293, Amph1, pMAP1S-S900 LC, and β-actin. (*C*) Quantification of the phosphorylation levels of Amph1-S293 normalised to total Amph1. Bars indicate mean ± SEM. ns, not significant by two-way ANOVA followed by Sidak’s multiple comparison test. WT *n* = 3 coverslips/condition, KO *n* = 3 coverslips/condition from 3 independent preparations of neuronal cultures. *(D)* EGCG, PD98059, and roscovitine were combined to block the kinase activity of Dyrk1A, MAPK, and Cdk5, respectively. All three kinases phosphorylate Amph1 in neurons with Dyrk1A and MAPK (continuous lines) and possibly Cdk5 (dashed line) also targeting S293. The skeletal structures of the inhibitors were generated by ACD/ChemSketch, 2021.1.0. *(E)* Hippocampal neurons at 14-15 DIV derived from WT and CDKL5 KO rats were treated with 20 mM EGCG, 100 μM PD98059, and 50 μM roscovitine inhibitors combined together for 1 h and stimulated with 50 mM KCl prior to repolarisation for 10 min in the presence of kinase inhibitors, or appropriate amount of DMSO. Lysates were assessed for pAmph1-S293, Amph1, pMAP1S-S900 LC, and β-actin. *(F)* Quantification of pAmph1-S293 levels normalised to total Amph1. Background was subtracted in all cases. Bars indicate mean ± SEM. ns, not significant by two-way ANOVA followed by Sidak’s multiple comparison test. WT *n* = 3 coverslips/condition, KO *n* = 3 coverslips/condition from 3 independent preparations of neuronal cultures.

Previous studies determined Amph1-S293 as the residue phosphorylated by CDKL5 *in vitro* (Katayama et al., 2015; Sekiguchi et al., 2013), which also resides within a CDKL5 consensus motif (Baltussen et al., 2018; Munoz et al., 2018). Furthermore, Amph1-S293 appears to be a plausible CDKL5 target in relation to its potential role in SV endocytosis, since its phosphorylation status regulates the affinity of Amph1 for the presynaptic endocytosis protein endophilin A1 (Murakami et al., 2006; Sekiguchi et al., 2013). To explore CDKL5-mediated phosphorylation of Amph1, we generated a rabbit polyclonal phospho-specific antibody against Amph1-S293 (**Figure S4A**). To validate this antibody, we generated recombinant GST-conjugated constructs of the central region of WT Amph1 that encompassed this site (residues 248-620, GST-Amph1) and two phospho-mutants, a null (GST-S293A) and a mimetic (GST-S293E) and assessed its specificity by Western blotting. This approach revealed that the pAmph1-S293 antibody reacted exclusively with the phospho-mimetic GST-S293E (**Figure S4B**), suggesting that the phospho-antibody is highly specific for phosphorylated Amph1-S293.

Amph1 undergoes dephosphorylation coupled to neuronal activity (Bauerfeind et al., 1997; Micheva et al., 1997). Accordingly, Amph1-S293 is one of the phospho-sites that is dephosphorylated following high frequency stimulation (Craft et al., 2008; Murakami et al.,2006). Therefore, we next focused on verifying whether Amph1-S293 was dephosphorylated in an activity-dependent manner. Initially, we treated hippocampal neuronal cultures with 50 mM KCl for 2 min to induce neuronal depolarisation. This greatly reduced the signal from the pAmph1-S293 antibody when compared to basal cultures, suggesting that the antibody accurately reports the phosphorylation status of this residue. We next examined whether Amph1-S293 dephosphorylation occurs via calcineurin, since this Ca^2+^-dependent enzyme dephosphorylates a series of presynaptic proteins during neuronal activity (Bauerfeind et al.,1997; Cousin and Robinson, 2001; Marks and McMahon, 1998; Nichols et al., 1994). Treatment with cyclosporin A, a calcineurin inhibitor, prevented the activity-dependent dephosphorylation at Amph1-S293, confirming that calcineurin performs this role. In contrast, treatment with calyculin A, an inhibitor of protein phosphatases 1 and 2A that are responsible for the main phosphatase activity in presynaptic terminals under basal and depolarising conditions, failed to prevent Amph1-S293 dephosphorylation (**Figure S4C**). Additionally, we examined the impact of electrical field stimulation, during which neurons were stimulated with 300 APs at 10 Hz or 400 APs at 40 Hz in the presence or absence of the antagonists AP5 and CNQX (which prevent postsynaptic activity or recurrent spontaneous activity). We observed that Amph1-S293 was dephosphorylated after stimulation at both frequencies (**Figure S4D**). Furthermore, the phosphorylation profile of Amph1-S293 was similar to that of pDyn1-S774, an established phosphorylation site that undergoes calcineurin- and activity-dependent dephosphorylation (Clayton et al., 2009; Liu et al., 1994; Tan et al.,2003). Overall, these findings suggest that Amph1-S293 undergoes calcineurin-mediated dephosphorylation linked to neuronal activity at the presynapse.

To assess whether Amph1-S293 is a CDKL5 substrate, WT and CDKL5 KO neuronal cultures were stimulated with 50 mM KCl and allowed to repolarise for different periods of increased duration to determine whether the absence of CDKL5 impacted on rephosphorylation of this residue (**Figure 7B**). KCl stimulation was employed to ensure complete dephosphorylation of S293, providing the widest possible dynamic range to visualise changes in its rephosphorylation. A phospho-antibody against the established endogenous CDKL5 substrate microtubule-associated protein 1S (MAP1S)-S900 was also used as a positive control (Baltussen et al., 2018; Munoz et al., 2018). In WT neurons, Amph1-S293 was efficiently rephosphorylated within 2.5 minutes after KCl stimulation (**Figure 7C**). In CDKL5 KO neurons there was no significant change in the phosphorylation levels of Amph1-S293 either before, during or after the KCl stimulus when compared to WT controls (**Figure 7C**). In contrast, phosphorylation of MAP1S-S900 was eliminated in CDKL5 KO neurons in all conditions. This supports the conclusion that Amph1-S293 is not directly phosphorylated by CDKL5 *in vivo* and, therefore, this phospho-site does not play a significant role in the slowing of SV endocytosis due to CDKL5 deficiency.

### Amph1-S293 is phosphorylated independently of CDKL5 at the presynapse

The unaltered phosphorylation levels of Amph1-S293 in the absence of CDKL5 indicates that another protein kinase is responsible for its phosphorylation *in vivo.* However, it is also possible that a different protein kinase substitutes for CDKL5 activity in CDKL5 KO neurons. A number of early studies showed that there are two protein kinases that phosphorylate Amph1-S293 *in vitro* in addition to CDKL5, including dual-specificity tyrosine phosphorylation-regulated kinase 1A (Dyrk1A) (Murakami et al., 2006) and mitogen-activated protein kinase (MAPK) (Shang et al., 2004), whereas cyclin-dependent kinase 5 (Cdk5) (Floyd et al., 2001; Liang et al., 2007) is also reported as an Amph1 kinase in mature neurons. In an attempt to unmask any potential phosphorylation of Amph1-S293 and to determine whether other protein kinases may substitute for CDKL5 in its absence, we treated WT and CDKL5 KO neurons with a cocktail of protein kinase inhibitors, including epigallocatechin gallate (EGCG), PD98059, and roscovitine to simultaneously eliminate the kinase activity of Dyrk1A, MAPK, and Cdk5, respectively (**Figure 7D**). KCl-induced depolarisation of WT and CDKL5 KO neurons was followed by repolarisation for 10 min (Error! Reference source not found.**Figure 7E**). We revealed that the phosphorylation levels of pAmph1-S293 were not altered between genotypes when normalised to total Amph1, as previously observed (**Figure 7F**). Moreover, the cocktail of kinase inhibitors abolished the rephosphorylation of pAmph1-S293 post-stimulation, indicating that kinases other than CDKL5 phosphorylate this residue in WT neurons and the contribution of CDKL5 to its phosphorylation is minor, if any. Importantly, the unaltered phosphorylation of the endogenous CDKL5 substrate pMAP1S-S900 (Baltussen et al., 2018; Munoz et al., 2018) in the presence of inhibitors excludes the possibility these inhibitors to act on CDKL5. Collectively, these data suggest that at least one presynaptic kinase other than CDKL5 phosphorylates pAmph1-S293 at hippocampal neurons.

## Discussion

CDD is emerging as a prominent monogenic neurodevelopmental and epileptic encephalopathy, therefore determining the key biological roles of CDKL5 will be vital in developing targeted therapies. In this work, we reveal the first direct role for CDKL5 at the presynapse, the control of SV regeneration. This requirement was specific to SV regeneration, with no other aspects of the SV life cycle impacted by the absence of the kinase. This defect in CDKL5 KO neurons was stimulus-independent, suggesting CDKL5 performs a fundamental role in the facilitation of this process. Importantly, CDKL5 protein kinase activity was both necessary and sufficient for this role, suggesting that CDKL5-dependent phosphorylation performs a fundamental role in facilitating SV turnover during neuronal activity.

A number of postsynaptic defects has been observed in a series of CDKL5 KO model systems, such as increased (Okuda et al., 2017; Yennawar et al., 2019) or decreased (Della Sala et al.,2016) long-term potentiation, altered dendritic morphology/dynamics (Amendola et al.,2014; Della Sala et al., 2016; Tang et al., 2017; Terzic et al., 2021), upregulated NMDA receptor number (Okuda et al., 2017; Tang et al., 2019; Terzic et al., 2021) and a shift in AMPA receptor subunit composition (Yennawar et al., 2019). Furthermore, a number of studies have suggested that loss of CDKL5 impacts synapse numbers in specific brain regions. Alterations in synapse number have been proposed to modulate the frequency of miniature events in systems where CDKL5 is absent (Della Sala et al., 2016; Ricciardi et al., 2012). However, in our primary neuronal culture system, we observe no obvious change in synapse number via staining with the presynaptic marker SV2A. Our study takes advantage of a novel rodent system to model CDD, a CDKL5 KO rat. A full characterisation of the electrophysiological and behavioural phenotypes of the CDKL5 rat model is described here (Simões de Oliveira et al. 2022), however similarly to other constitutive CDKL5 KO models, they do not display overt seizure activity. Hippocampal brain slices from this model system do display reduced mEPSC frequency with no apparent decrease in synapse number (Simões de Oliveira et al. 2022), suggesting that this defect may be linked to dysfunctional SV regeneration rather than less available synapses.

We revealed that the kinase activity of CDKL5 is necessary and sufficient for its role in SV regeneration. This was achieved via use of structural/patient mutations and expression of isolated domains in molecular replacement studies. The K42R mutant is a *bona fide* kinase dead protein since it fails to bind ATP and phosphorylate targets *in vitro* (Lin et al., 2005). The patient mutation R178P (Elia et al., 2008; Nemos et al., 2009) is also assumed to be kinase dead since a similar patient mutation (R178W) abolished kinase activity *in vitro* (Munoz et al.,2018). However the kinase activity of this specific mutant still has to be directly investigated. The ability of the isolated CDKL5 kinase domain to fully restore presynaptic function was surprising, and suggests that the unstructured C-terminus, which forms the majority of the protein, is dispensable for CDKL5 localisation and/or substrate recognition. Importantly, overexpression of full-length protein did not affect SV regeneration, suggesting increased gene dosage is not deleterious to presynaptic function. These findings have important implications for future gene therapy studies, since insertion of the isolated kinase domain will facilitate packaging inside viral delivery vectors that have limited space. The potential of this strategy to have therapeutic benefits in individuals with CDD is supported by studies where re-expression of the CDKL5 gene in KO mice fully reversed a cohort of cell, circuit and behavioural phenotypes (Terzic et al., 2021). The finding that CDD appears to be a disorder of neuromaintenance and not neurodevelopment (Kind and Bird, 2021), provides support that expression of the CDKL5 kinase domain later in life may restore specific aspects of brain function.

We also revealed that S293 on Amph1 is not the CDKL5 substrate that controls SV regeneration. This site was an excellent candidate, since it was situated within a CDKL5 consensus sequence, is phosphorylated by the kinase *in vitro* (Katayama et al., 2015; Sekiguchi et al., 2013) and is the dominant *in vivo* site on Amph1 (Craft et al., 2008). Furthermore, this site is dephosphorylated during neuronal activity (Craft et al., 2008; Murakami et al., 2006) and its phosphorylation status controls interactions with the endocytosis protein endophilin (Murakami et al., 2006; Sekiguchi et al., 2013). Finally, deficiency of Amph1 results in occurrence of irreversible seizures in mice (Di Paolo et al., 2002). However, phospho-specific antibodies against S293 revealed no change in its phosphorylation status in CDKL5 KO neurons. A series of *in vitro* studies have identified other candidate proteins kinases that could phosphorylate this site (Floyd et al., 2001; Liang et al., 2007; Murakami et al., 2006; Shang et al., 2004). An inhibitor cocktail containing antagonists of these protein kinases abolished rephosphorylation of Amph1 S293 in both WT and CDKL5 KO neurons, suggesting that these protein kinases do not substitute for CDKL5 in its absence. Given the interplay between CDKL5 and Dyrk1A (Oi et al., 2017; Trovo et al., 2020), we also excluded the possibility that CDKL5 loss may influence the phosphorylation of Amph1-S293 by Dyrk1A. The identity of the protein kinase that rephosphorylates S293 is still therefore undetermined, however it is clear that its phosphorylation does not mediate CDKL5-dependent effects on SV regeneration. The identity of the presynaptic CDKL5 substrate(s) is currently under investigation.

One interesting observation was that the impact of loss of CDKL5 function on SV regeneration appeared to reduce with increasing stimulus frequencies. Remarkably, CDKL5 is not the only kinase of the CMGC (named after the initials of some member kinases) group that has been reported to behave in a frequency-dependent manner. For example, overexpression of Dyrk1A results in more profound SV endocytosis delay following low rather than high frequencies in hippocampal neurons (Kim et al., 2010). This is an intriguing observation, since defects in SV endocytosis are typically exacerbated with increased stimulus intensities (McAdam et al., 2020; Zhao et al., 2014). Since GABAergic neurons usually fire at higher frequencies (Bartos et al., 2007), this suggests that excitatory neurotransmission may be disproportionately affected by the absence of CDKL5. Recent studies in conditional CDKL5 KO models provide some support to this hypothesis. For example, selective deletion of CDKL5 in inhibitory interneurons increases mEPSC, but not mIPSC frequency (Tang et al., 2019). Furthermore, conditional KO of CDKL5 in mouse excitatory neurons resulted in overt seizure phenotypes (with increased mEPSCs, but not mIPSCs), whereas the equivalent deletion in inhibitory neurons had little effect (Wang et al., 2021). Therefore there appears to be a complex relationship between loss of CDKL5 function when examined at the level of intact brain circuits. Consequently, it may be too soon to predict how defects in presynaptic SV regeneration culminate in both global and specific circuit dysfunction and ultimately seizure activity in individuals with CDD.

In summary, we have identified a key presynaptic role for CDKL5 in neurotransmission and potentially circuit and brain function. It will be critical to determine the molecular target(s) of this kinase within this specialised subcellular region to determine the extent that presynaptic dysfunction underpins this neurodevelopmental and epileptic encephalopathy.

## Acknowledgements

This work was supported by grants from the Loulou Foundation, the Simons Foundation (529508), The Wellcome Trust (204954/Z/16/Z) and a College of Medicine and Veterinary Medicine studentship to CK. For the purpose of open access; the author has applied a CC-BY public copyright license to any author accepted manuscript version arising from this submission. We thank Lynsey Dunsmore for help with maintenance of the *Cdkl5* KO LE colony and Dr. Sally Till for her help with the cortices dissection. We also thank Dr Ira Milosevic and Dr Giles Hardingham for useful suggestions in an earlier version of this work.

## Author Contributions

Conceptualization, PCK, MAC; Methodology CK, ECD, MAC; Data analysis, CK; Visualisation, CK; Investigation, CK, MAC; Resources, PCK; Writing, CK, MAC; Funding Acquisition, PCK, MAC.

## Star Methods

**Key Resources Table.**

| REAGENT or RESOURCE | SOURCE | IDENTIFIER |
| --- | --- | --- |
| **Antibodies** |  |  |
| Chicken anti-GFP | Abcam | Cat# ab13970, RRID:AB_300798 |
| Donkey anti-goat IRDye 680RD | LI-COR Biosciences | Cat# 925-68074, RRID:AB_2650427 |
| Donkey anti-goat IRDye 800CW | LI-COR Biosciences | Cat# 925-32214, RRID:AB_2687553 |
| Donkey anti-guinea pig IRDye 680RD | LI-COR Biosciences | Cat# 925-68077, RRID:AB_2814914 |
| Donkey anti-mouse IRDye 680RD | LI-COR Biosciences | Cat# 925-68072, RRID:AB_2814912 |
| Donkey anti-mouse IRDye 800CW | LI-COR Biosciences | Cat# 925-32212, RRID:AB_2716622 |
| Donkey anti-rabbit Alexa Fluor 488 | Molecular Probes, Thermo Fisher Scientific | Cat# A-21206, RRID:AB_2535792 |
| Donkey anti-rabbit IRDye 680RD | LI-COR Biosciences | Cat# 925-68073, RRID:AB_2716687 |
| Donkey anti-rabbit IRDye 800CW | LI-COR Biosciences | Cat# 925-32213, RRID:AB_2715510 |
| Donkey anti-sheep Alexa Fluor 568 | Molecular Probes, Thermo Fisher Scientific | Cat# A-21099, RRID:AB_2535753 |
| Goat anti-Amph1 | Santa Cruz Biotechnology | Cat# sc-8536, RRID:AB_2226798 |
| Goat anti-CHC | Santa Cruz Biotechnology | Cat# sc-6579, RRID:AB_2083170 |
| Goat anti-chicken Alexa Fluor 488 | Molecular Probes, Thermo Fisher Scientific | Cat# A-11039, RRID:AB_142924 |
| Goat anti-Dyn1 | Santa Cruz Biotechnology | Cat# sc-6402, RRID:AB_639941 |
| Goat anti-EHD | Santa Cruz Biotechnology | Cat# sc-23452, RRID:AB_2097347 |
| Goat anti-guinea pig Alexa Fluor 568 | Molecular Probes, Thermo Fisher Scientific | Cat# A-11075, RRID:AB_141954 |
| Goat anti-syndapin 1 | Santa Cruz Biotechnology | Cat# sc-10412, RRID:AB_653788 |
| Guinea pig anti-VGLUT1 | Synaptic Systems | Cat# 135 304, RRID:AB_887878 |
| Mouse anti-PSD95 | BioLegend | Cat# 810401, RRID:AB_2564750 |
| Mouse anti-β-actin-peroxidase | Sigma-Aldrich | Cat# A3854, RRID:AB_262011 |
| Rabbit anti-ATP6V1B2 | Abcam | Cat# ab183887 |
| Rabbit anti-CDKL5 | Atlas Antibodies | Cat#HPA002847, RRID:AB_1080107 |
| Rabbit anti-pAkt-S473 | Cell signalling | Cat# 9018, RRID:AB_2629283 |
| Rabbit anti-pAmph1-S293 | MRC Protein Phosphorylation Unit, University of Dundee, UK | N/A |
| Rabbit anti-pGSK3α/β-S9/S21 | Cell signalling | Cat# 9331, RRID:AB_329830 |
| Rabbit anti-pMAP1S (light chain)-S900 | Rouse lab, University of Dundee, UK | N/A |
| Rabbit anti-sheep IRDye800 conjugated | Rockland Immunochemicals | Cat# 613-732-168, RRID:AB_220182 |
| Rabbit anti-SV2A | Abcam | Cat# ab32942, RRID:AB_778192 |
| Rabbit anti-Syp1 | Abcam | Cat# ab14692, RRID:AB_301417 |
| Sheep anti-CDKL5 (Human epitope) | Rouse lab, University of Dundee, UK | N/A |
| Sheep anti-CDKL5 (Mouse epitope) | Rouse lab, University of Dundee, UK | N/A |
| Sheep anti-pDyn1-S774 | AbD Serotec (Bio-rad) | Cat# AHP899, RRID:AB_567392 |
| **Chemicals, peptides, and recombinant proteins** |  |  |
| B-27™ Supplement (50X), serum free | Gibco™, ThermoFisher Scientific | Cat# 17504044 |
| Bafilomycin A1 | Alfa Aesar | Cat# J61835.MX |
| Boric Acid | Sigma-Aldrich | Cat# B6768 |
| Bovine serum albumin | Roche Diagnostics GmbH | Cat# 10735078001 |
| Bradford | AppliChem | Cat# A6932 |
| Calyculin A | Abcam | Cat# ab141784 |
| Cyano-7-nitroquinoxaline-2,3-dione | Abcam | Cat# ab120271 |
| Cyclosporin A | Sigma-Aldrich | Cat# 30024 |
| Cytosine arabinofuranoside | Sigma-Aldrich | Cat# C1768 |
| Dimethyl sulfoxide | Sigma-Aldrich | Cat# D8418 |
| Dithiothreitol | Sigma-Aldrich | Cat# D0632 |
| DL-2-Amino-5-phosphonopentanoic acid | Abcam | Cat# ab120044 |
| DMEM-F12 | Gibco | Cat# 21331-020 |
| Epigallocatechin gallate | Calbiochem | Cat# 324880-10MG |
| Foetal Bovine serum | BioSera | Cat# S1810-500 |
| Glutathione Sepharose 4B beads | GE Healthcare | Cat# GE17-0756-01 |
| Instant Blue™ Protein Stain | C.B.S. Scientific | Cat# HG73010 |
| Intercept (PBS) Blocking Buffer | LI-COR Biosciences | Cat# 927-70001 |
| Intercept (TBS) Blocking Buffer | LI-COR Biosciences | Cat# 927-60001 |
| Isopropyl β-D-1-thiogalactopyranoside | Calbiochem | Cat# 420322 |
| Laminin | Sigma-Aldrich | Cat# L2020 |
| L-glutamine | Gibco | Cat# 25030-024 |
| Lipofectamine 2000 | ThermoFisher Scientific | Cat# 11668027 |
| Lysozyme | Sigma-Aldrich | Cat# L6876 |
| Neurobasal | Gibco | Cat# 21103-049 |
| Papain | Worthington Biochemical Corporation | Cat# LK003178 |
| PD98059 | EMD Millipore Corp., | Cat# 513000-5MG |
| Penicillin/Streptomycin | Gibco | Cat# 15140-122 |
| Poly-D-lysine | Sigma-Aldrich | Cat# P7886 |
| Ponceau-S staining | Sigma-Aldrich | Cat# P7170 |
| ProbeQuant G-50 Micro Column | GE Healthcare | Cat# 28903408 |
| Protease inhibitor cocktail | Sigma-Aldrich | Cat# P8849 |
| Protein G Agarose beads | Sigma-Aldrich | Cat# 11719416001 |
| Roscovitine | EMD Millipore Corp. | Cat# 557360-1MG |
| **Experimental Models: organisms/strains** |  |  |
| Mouse: C57Bl/6J | Charles River, UK | N/A |
| Rat: *Cdkl5* KO LE, Long-Evans | Horizon Discovery, USA | N/A |
| Rat: Sprague Dawley | Charles River, UK | N/A |
| **Recombinant DNA** |  |  |
| GST-Amph1 | This paper | N/A |
| GST-Amph1 S293A | This paper | N/A |
| GST-Amph1 S293E | This paper | N/A |
| Human CDKL5 isoform 1 (CDKL5) | Dr. V. Kalscheuer, Max Planck Institute for Molecular Genetics, DE | N/A |
| mCerC1 | (Gordon and Cousin, 2013) | N/A |
| mCer-hCDKL5 isoform 1 | This paper | N/A |
| mCer-hCDKL5 K42R isoform 1 | This paper | N/A |
| mCer-hCDKL5 R178P isoform 1 | This paper | N/A |
| mCer-hCDKL5 ΔC isoform 1 | This paper | N/A |
| mCer-hCDKL5 Δkinase isoform 1 | This paper | N/A |
| pGEX-KG | Dr. C. Rickman, Heriot-Watt University, UK | N/A |
| Rat Amph1 | Dr. H. T. McMahon, MRC Laboratory of Molecular Biology, UK | N/A |
| Syp1-mCer | (Gordon et al., 2011) | N/A |
| sypHy | Prof. L. Lagnado, University of Sussex | RRID:Addgene_24478 |
| **Software and Algorithms** |  |  |
| ACD/ChemSketch 2021.1.0 | ACD/Labs | RRID:SCR_019272 |
| Adobe Illustrator | Adobe Inc. | RRID:SCR_010279 |
| Fiji ImageJ | https://imagej.net/software/fiji/ | RRID:SCR_002285 |
| GraphPad Prism 8.4.2 | GraphPad Software Inc. | RRID:SCR_002798 |
| Image Studio Lite Ver 5.2 | LI-COR Biosciences | RRID:SCR_013715 |

### Resource Availability

#### Lead Contact

Further information and requests for resources and reagents should be directed to and will be fulfilled by the Lead Contact, Michael Cousin.

#### Materials Availability

All unique/stable reagents generated in this study are available from the Lead Contact without restriction.

#### Data and Code Availability

This study did not generate datasets/code.

## EXPERIMENTAL MODEL AND SUBJECT DETAILS

### Rats

All experimental procedures were conducted according to the UK Animal (Scientific Procedures) Act 1986 on the protection of animals used for scientific purposes and were approved by the Animal Welfare and Ethical Review Body at the University of Edinburgh (Home Office project license to M. Cousin – 7008878 or D. Wyllie - P1351480E). Adult animals were killed by exposure to increasing CO_2_ concentration followed by cervical dislocation, while embryos were killed by decapitation followed by destruction of the brain. All animals were maintained on a 12-hour light/dark cycle under constant temperature, with food and water provided when needed.

*Cdkl5* KO Long-Evans rats were generated by Horizon Discovery, USA, following a CRISPR interference approach to delete 10 bp in exon 8 of the *Cdkl5* gene (138367-76 in genomic sequence) that results in the introduction of an early stop codon (Simões de Oliveira et al. 2022). *Cdkl5* heterozygous females (*Cdkl5*^+/−^) were crossed with WT Long-Evans males (*Cdkl5*^+/y^) and the offspring were obtained from pregnant females at E17-E19. Prior to genotyping, embryos were sexed by dissecting the abdomen to reveal their inner reproductive organs. Male *Cdkl5^−/y^* embryos (referred to as CDKL5 KO) and male WT littermate controls were used for neuronal cultures. WT and CDKL5 KO adult (> 2 months old) male rats were used for biochemistry experiments. For CV analysis, primary hippocampal cultures were prepared from WT mouse embryos (C57BL/6J) at E16-18.

### Genotyping

Genomic DNA was obtained from nose or tail biopsies of embryos with alkaline reagent containing 25 mM NaOH and 0.2 mM disodium EDTA (pH 12) at 95 °C (HotSHOT). DNA extract (1 μl) was used for genotyping with the following primers (Eurogentec, BE): 5’-GGGCTTGTAGCAAATCCATCC-3’ (sense), 5’-ATACGTGGCTACTCGGTGGTAC-3’ (sense; matching 10 bp deletion), and 5’-AGCAAGCAGAGTTCTATTTTCCT-3’ (antisense) using polymerase chain reaction.

## METHOD DETAILS

### DNA constructs

The plasmid DNA vectors in this study were obtained as follows: sypHy from Prof. L. Lagnado (University of Sussex, UK), full-length human CDKL5_1 (hCDKL5_1; referred to as CDKL5) from Dr. V. Kalscheuer (Max Planck Institute for Molecular Genetics, Berlin, Germany), full-length rat Amph1 from Dr. H. T. McMahon (MRC Laboratory of Molecular Biology, Cambridge, UK), and pGEX-KG from Dr. C. Rickman (Heriot-Watt University, Edinburgh, UK).

mCerulean (mCer)-C1-CDKL5 was generated by subcloning CDKL5 into an mCerC1 vector, where the original GFP moiety was replaced by mCer (Gordon and Cousin, 2013), with the primers 5’-CATCAT<u>CTCGAG</u>GAATGAAGATTCCTAACATTGGTAATG-3’ (sense) and 5’-CATCAT<u>GGTACC</u>TTACAAGGCTGTCTCTl llAAATC-3’ (antisense) with restriction sites underlined. Deletion mutants of CDKL5 were generated using the subsequent primers: 5’-CATCAT<u>CTCGAG</u>TAATGAAGATTCCTAACATTGG-3’ (sense) and 5’-ATGATG<u>GAATTC</u>CTAAAATGTAGGGTGATTCAAAC-3’ (antisense) for the kinase domain (residues 1-297) and 5’-CATCAT<u>CTCGAG</u>TACAAACCCAGAGACTTCTGG-3’ (sense) and 5’-ATGATG<u>GGTACC</u>TTACAAGGCTGTCTCTTTTAAATC-3’ (antisense) for the C-terminal tail (residues 298-960) with restriction sites underlined. Point mutations were introduced into CDKL5 using standard site-directed mutagenesis protocols with the following primers: 5’-GAAATTGTGGCGATC<u>CG</u>GAAATTCAAGGACAGT-3’ (sense) and 5’-ACTGTCCTTGAATTTC<u>CG</u>GATCGCCACAATTTC-3’ (antisense) for K42R and 5’-GCCACCAGATGGTATC<u>C</u>GTCCCCAGAACTCTTA-3’ (sense) and 5’-TAAGAGTTCTGGGGAC<u>G</u>GATACCATCTGGTGGC-3’ (antisense) for R178P with mutated sites underlined. GST-Amph1 was generated by subcloning Amph1 (residues 248-620) into a pGEX-KG vector using the primers 5’-CATCAT<u>GAATTC</u>TAGGAGCTCCCAGTGATTCGGGTC-3’ (sense) and 5’-ATGATG<u>CTCGAG</u>CTAAGGAGGCAGTTCCTGAGCGG-3’ (antisense) with restriction sites underlined. Point mutations were introduced into Amph1 using standard site-directed mutagenesis protocols with the following primers: 5’-CCAGTGCGACCCAGA<u>G</u>CACCTTCACAGACAAGG-3’ (sense) and 5’-CCTTGTCTGTGAAGGTG<u>C</u>TCTGGGTCGCACTGG-3’ (antisense) for S293A and 5’-CCAGTGCGACCCAGA<u>GA</u>ACCTTCACAGACAAGG-3’ (sense) and 5’-CCTTGTCTGTGAAGGT<u>TC</u>TCTGGGTCGCACTGG-3’ (antisense) for S293E with mutated sites underlined. All constructs were validated by Sanger sequencing.

### Neuronal cultures and transfection

Hippocampi were dissected from CDKL5 KO male embryos and littermate controls and dissociated in papain (10.5 U/ml). Tissue was triturated in Dulbecco’s Modified Eagle Medium/Nutrient Mixture F-12 supplemented with 10 % (v/v) foetal bovine serum. Following a low-speed centrifugation, neurons were resuspended in Neurobasal medium supplemented with 0.5 mM L-glutamine, 1 % (v/v) B27 supplement, and penicillin/streptomycin. Neurons were plated on poly-D-lysine- and laminin-precoated coverslips and kept in supplemented Neurobasal medium in a humidified incubator at 37 °C/5 % CO_2_ for up to 15 days. Cytosine β-D-arabinofuranoside was added to neurons at 1 μM on 3 DIV to prevent glial proliferation. Neurons were transfected after 8-9 DIV with Lipofectamine 2000 as per manufacturer’s instructions.

### Live-cell imaging and data analysis

Primary hippocampal neurons at 13-15 DIV were mounted in a closed bath imaging chamber (RC-21BRFS, Warner) allowing electrical field stimulation (1-ms pulse width, 100 mA current output). Tyrode’s buffer (119 mM NaCl, 2.5 mM KCl, 2 mM CaCl_2_, 2 mM MgCl_2_, 25 mM HEPES, 30 mM glucose, pH 7.4), supplemented with 10 μM 6-cyano-7-nitroquinoxaline-2,3-dione (CNQX) and 50 μM DL-2-amino-5-phosphonopentanoic acid (AP5) was perfused continuously. At the end of each recording, neurons were perfused with 50 mM NH_4_Cl solution, pH 7.4, substituting equal concentration of NaCl in Tyrode’s buffer. All recordings were performed at room temperature. Transfected neurons were visualized using a Zeiss Axio Observer D1 inverted epifluorescence microscope (Zeiss Ltd., Germany) with a 40x 1.3 NA oil immersion objective. Time-lapse images were acquired using a Hamamatsu Orca-ER camera and the acquisition rate was set at 4 s constantly pre- and post-stimulation. Neurons expressing sypHy and mCer constructs were imaged at 500 nm and 430 nm excitation, respectively, using a 525-nm dichroic and a 535-nm emission filter. To measure exocytosis rate, Tyrode’s buffer was supplement with 1 μM bafilomycin A1. To measure acidification kinetics, HEPES was replaced by 25 mM 2-(N-morpholino)ethanesulfonic acid in Tyrode’s buffer and acquisition rate was set at 2 s.

Time-lapse stacks of images were analysed using the Fiji is just ImageJ (Fiji) software (Schindelin et al., 2012; Schneider et al., 2012). These were initially aligned using the StackReg plugin with Rigid Body transformation type (Thevenaz et al., 1998). Regions of interest of 0.8 μm in diameter were placed on presynaptic boutons responsive to stimulation. Fluorescence intensity was measured for all image slices using the Times Series Analyzer (https://bit.ly/3M5hWpb). The average ΔF/F0 was calculated for each coverslip were normalised to the maximum fluorescence intensity either during stimulation or NH4Cl perfusion. A one-phase exponential fit was used to correct baseline for bleaching that was subtracted from all time points. Distance to the baseline at fixed time (122 s after termination of stimulation) was used as a measure of endocytosis speed. No background was subtracted.

### Immunocytochemistry

Primary cultured hippocampal neurons were fixed with 4 % (w/v) paraformaldehyde/PBS for 10 min and neutralized with 50 mM NH4Cl/PBS for 10 min. After washing with PBS, neurons were permeabilized in 0.1 % (v/v) Triton X-100, 1 % (w/v) bovine serum albumin/PBS for 5 min and blocked in 1 % (w/v) bovine serum albumin/PBS for 30 min. Following blocking, neurons were incubated with the appropriate dilution of primary antibodies for 1-2 h at room temperature. Primary antibodies were used as follows: sheep CDKL5 (1:200), chicken GFP (1:5000), rabbit SV2A (1:200), and guinea pig VGLUT1 (1:1000). Alexa Fluor secondary antibodies (1:1000) were applied for 1-2 h at room temperature in the dark.

Transfected neurons were visualized using a Zeiss Axio Observer Z1 inverted epifluorescence microscope (Zeiss Ltd., Germany) and a 40x 1.3 NA oil immersion objective at 480 nm and 550 nm excitation wavelengths. Fluorescent light was detected at 500-552 nm and >565 nm using a 495-nm and a 565-nm dichroic filter, respectively. Neurons expressing mCer-tagged constructs were visualized at 480 nm excitation wavelength using the anti-GFP antibody described above. Images were acquired using a Zeiss AxioCam 506 camera and Zeiss ZEN 2 software. Data analysis was performed using Fiji. To quantify endogenous CDKL5 expression, regions of interest were drawn manually around GFP-expressing cell bodies and average CDKL5 signal was calculated and normalised to that of untransfected cell bodies. Background was subtracted in all cases. For counting bouton numbers, MaxEntropy thresholding was applied and positive accumulations of 0.64-2.24 μm in diameter were counted using the Analyze particles plugin (Kapur et al., 1985). The number of SV2A- and VGLUT1-positive puncta was counted in (50 × 15) μm^2^ selections along neuronal processes to eliminate the influence of neuronal density variation between genotypes. For CV analysis, the mean GFP fluorescence along an axonal segment of > 15 μm was divided by the standard deviation and expressed as a percentage (Gordon and Cousin, 2013). The average CV value of five axonal segments was calculated per field of view.

### Biochemical isolation of crude SVs

The crude purification of SVs was performed as described previously (Huttner et al., 1983). An adult rat brain was homogenized in ice-cold 0.32 M sucrose, 5 mM EDTA (pH 7.4) after removing the cerebellum. The homogenate (H) was centrifuged twice at 950 x *g* for 10 min at 4 °C and the supernatant was collected each time. The combined supernatant (S1) was spun at 20,400 x *g* for 30 min at 4 °C. The pellet (P2) represents the crude synaptosomal fraction. For crude isolation of SVs, the P2 fraction was resuspended in ice-cold 0.32 M sucrose/EDTA and incubated with 1 M HEPES/NaOH solution (pH 7.4) on ice for 30 min. After spinning at 32,900 x *g* for 20 min at 4 °C, the lysate pellet (LP1) and lysate supernatant (LS1) were obtained. The supernatant was then centrifuged at 268,000 x *g* for 2 h at 4 °C to generate LS2 and LP2 fractions. The LP2 pellet that represents the crude SV fraction was collected and resuspended in 40 mM sucrose. Aliquots of the intermediate fractions were kept for analysis. The total protein amount of the samples was measured by Bradford and their concentration was adjusted to 1 mg/ml prior to Western blot analysis.

### Immunoprecipitation

Adult rat brain was mechanically homogenized in buffer containing 50 mM HEPES (pH 7.5), 0.5 % (v/v) Triton X-100, 150 mM NaCl, 1 mM EDTA, 1 mM EGTA, 1 mM phenylmethylsulfonyl fluoride, and protease inhibitor cocktail. The homogenate was incubated for 1-2 h at 4 °C rotating and then centrifuged at 155,000 x *g* for 40 min at 4 °C. The supernatant was collected and pre-cleaned with Protein G Agarose beads (Sigma-Aldrich) for 1-2 h at 4 °C rotating to enhance specificity and the total protein content was quantified by Bradford assay. The brain lysate (equivalent to 2 mg of protein) was incubated with 2-4 μg of the antibody of interest at 4 °C rotating overnight. Next, Protein G Agarose beads (approximately 20 μl) were added to the antibody-containing brain lysates and left rotating for 1-2 h at 4 °C prior to being centrifuged at low speed. The supernatant was then discarded and after three washes in HEPES buffer, Laemmli sample buffer was added directly to the beads followed by heating at 95 °C for 5 min. A random antibody against Eps15 Homology Domain protein (EHD) was used as a control.

### Drug treatments

Cyclosporin A, calyculin A, PD98059 and roscovitine were dissolved in dimethyl sulfoxide (DMSO), whereas AP5, CNQX, PD98059 and EGCG in ultrapure water for stock concentration. For all drug experiments, culture medium was replaced by unsupplemented Neurobasal medium and neurons were treated with appropriate drug dilution at 37 °C. The drugs were administered as follows: 10 μM cyclosporin A or 100 nM calyculin A for phosphatase inhibition experiments, 50 μM AP5 and 10 μM CNQX for electrical stimulation, 20 mM EGCG, 100 μM PD98059, and 50 μM roscovitine for kinase inhibition experiments. Stimulation was performed at room temperature in the presence of drugs in Tyrode’s buffer prior to lysis with Laemmli buffer.

### Pull-down

Glutathione S-transferase (GST)-fused proteins were expressed in *Escherichia coli* BL21(DE3) cells in lysogeny broth medium containing ampicillin after induction with 1 mM isopropyl β-D-1-thiogalactopyranoside. Induced bacterial cultures were spun at 5000 x *g* for 15 min at 4 °C and the pellets were resuspended in ice-cold buffer containing 10 mM Tris, 150 mM NaCl, 1 mM EDTA, pH 8, protease inhibitors and 1 mM phenylmethylsulfonyl fluoride. Lysozyme (0.0675 μg/μl), 4 mM dithiothreitol, and 10 % (v/v) Triton X-100 were also added. The cells were sonicated at 10 kHz and the clear lysates were spun at 17,420 x *g* for 10 min at 4 °C. The supernatant was transferred to pre-washed Glutathione Sepharose 4B beads resuspended in PBS to create a 50 % suspension and left rotating overnight at 4 °C. A small volume of glutathione -coupled GST-fused proteins was loaded into a ProbeQuant G-50 Micro Column and washed once in ice cold lysis buffer containing 1 % (v/v) Triton X-100, 25 mM Tris-HCl, 150 mM NaCl, 1 mM EGTA, 1 mM EDTA, pH 7.4, prior to incubation with synaptosomal lysates. The columns were washed successively in ice cold lysis buffer, in NaCl-supplemented lysis buffer (500 mM) and in 20 mM Tris, pH 7.4. Laemmli sample buffer was added into the columns and the eluted proteins were denatured at 95 °C for 5 min. All GST-coupled Amph1 constructs were devoid of the N-terminal Bin/Amphiphysin/Rvs (N-BAR) and the Src-homology 3 (SH3) domains and their total level was estimated with Coomassie Brilliant blue staining prior to Western blot analysis.

### Western blotting

Brain samples were prepared as described above, whereas hippocampal neurons at 14-15 DIV were lysed directly with Laemmli sample buffer. Proteins were denatured at 95 °C for 5 min. Protein extracts were separated by SDS-PAGE, transferred to a nitrocellulose membrane, and blocked in Intercept (PBS or TBS) blocking buffer. Membranes were incubated with primary antibodies at 4 °C overnight and IRDye secondary antibodies (1:10000) for 2 h at room temperature in Intercept (PBS or TBS) blocking buffer containing 0.1 % (v/v) Tween-20 in the dark. Blots were visualized using the LI-COR Biosciences Odyssey Infrared Imaging System and quantification of band densities was performed using the Image Studio Lite version 5.2 software with background subtraction or Fiji. Equal protein amount loading was verified by Ponceau-S staining. The primary (phospho)antibodies that were used in this study are: sheep CDKL5 (1:500), goat Amph1 (1:500), goat CHC (1:250), goat syndapin 1 (1:1000), guinea pig VGLUT1 (1:2000), rabbit ATP6V1B2 (1:5000), goat Dyn1 (1:500), rabbit Syp1 (1:500), mouse PSD95 (1:1000), mouse β-actin-peroxidase (1:30000), rabbit pMAP1S-S900 (light chain) (1:50), sheep pDyn1-S774 (1:1000), rabbit pAkt-S473 (1:1000), rabbit pGSK3α/β-S9/S21 (1:1000). For experiments assessing the phosphorylation levels of Amph1-S293, a rabbit polyclonal phosphoantibody was raised against the peptide PVRPRS^293^PSQTRC of Amph1 (0.5 mg/ml).

### Statistical analysis and experimental design

Statistical calculations were conducted using GraphPad Prism 8.4.2 (GraphPad Software Inc). The normality of the data distribution was assessed by performing D’Agostino and Pearson omnibus normality test with significance level set at α= 0.05. Datasets following a Gaussian distribution were presented as mean ± standard error of the mean (SEM) and statistical significance was assessed by two-tailed unpaired t test for comparison between two groups or analysis of variance (ANOVA) followed by Tukey’s, Dunnett’s or Sidak’s post hoc analysis for multiple comparisons. Datasets following a non-Gaussian distribution were presented as median with interquartile range (IQR) indicating min to max whiskers and statistical significance was evaluated by Mann-Whitney test for comparison between two groups or Kruskal-Wallis followed by Dunn’s post hoc analysis for multiple comparisons. For experiments with a small number of replicates for a normality test to be performed, a parametric test was assumed. Asterisks refer to *p*-values as follows: *; *p* ≤ 0.05, **; *p* ≤ 0.005, ***; *p* ≤ 0.001, ****; *p* ≤ 0.0001. All experiments consisted of at least three independent biological replicates. Live-imaging data were analysed blind for experiments consisting of two groups. Random variation or effect size were not estimated. Sample size and statistical test are indicated in the figure legends. Detailed description of the statistical tests and p values are presented in **Table S1**.

