## Supplementary Data for "Epilepsy-related CDKL5 deficiency slows synaptic vesicle endocytosis in central nerve terminals"

### Supplementary figures

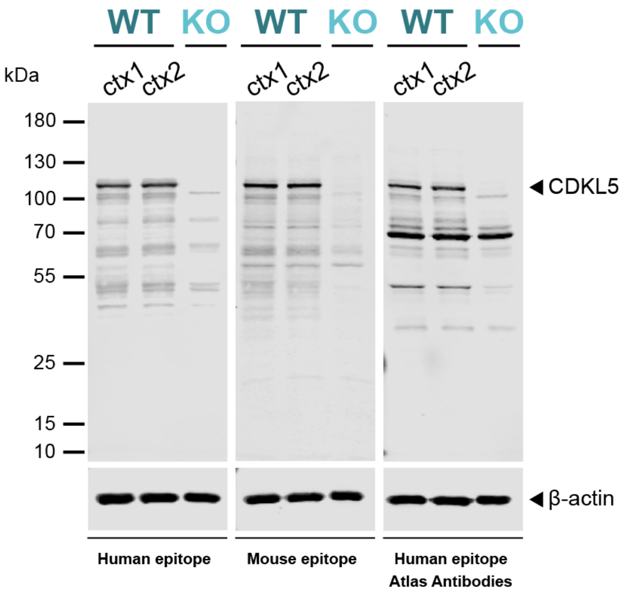


**Figure S1.** CDKL5 protein is absent from *Cdkl5* KO LE rats. Immunoblots of cortical lysates generated from WT and CDKL5 KO animals at P14 using three different antibodies against CDKL5. Two antibodies were raised against a human (aa 350-650) and a mouse (aa 300-600) epitope on CDKL5, respectively, whereas the Atlas CDKL5 antibody was tested as a commercially available alternative. CDKL5 is detected as a 110-kDa band that is absent from KO tissues. All three antibodies detect numerous non-specific bands observed in both WT and CDKL5 KO tissues. In all cases, β-actin was used as a loading control.

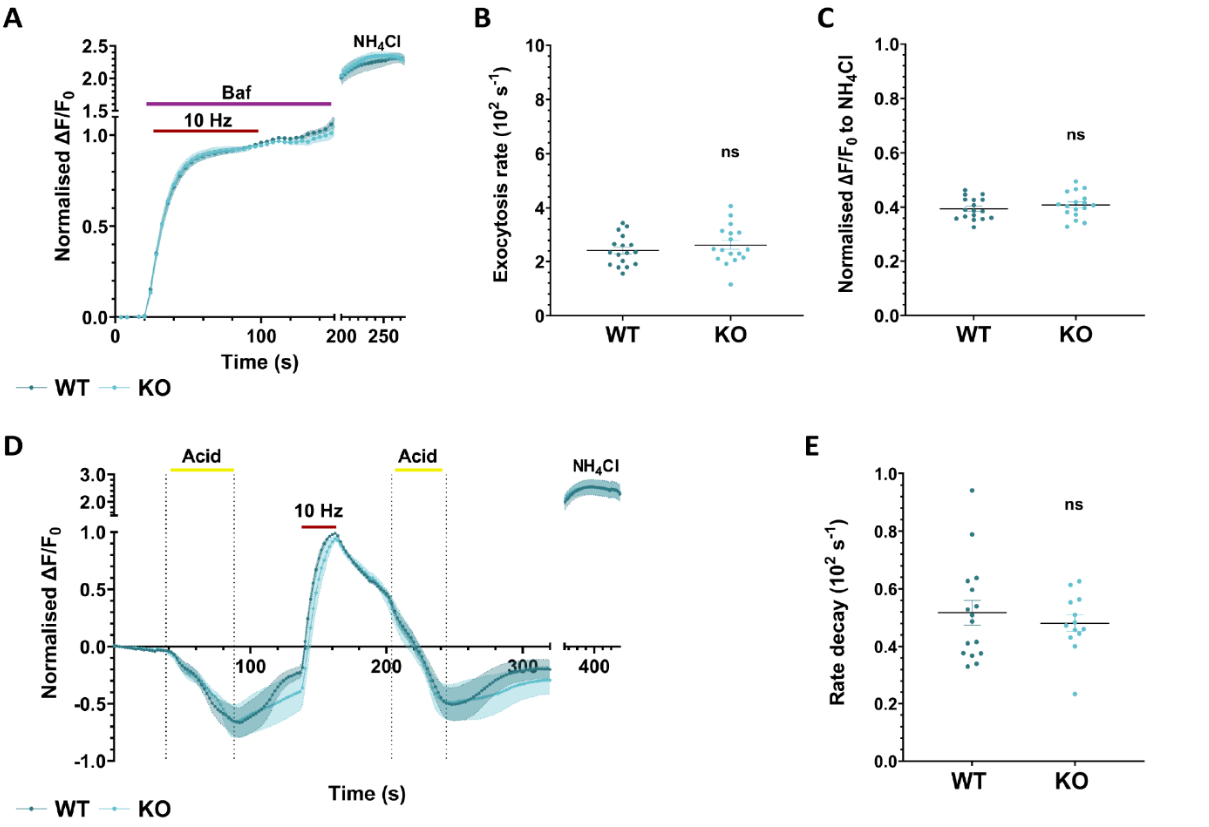


**Figure S2.** Loss of CDKL5 does not alter SV exocytosis rate, total pool size, or SV acidification rate. Primary hippocampal neurons from WT and CDKL5 KO rats were transfected with sypHy at 8-9 DIV and used at 13-14 DIV. (*A*) sypHy response from neurons stimulated with 900 APs at 10 Hz (red bar) in the presence of 1 µM bafilomycin A1 (purple bar) normalised to the plateau. (*B*) Quantification of exocytosis rate. Scatter plots indicate mean ± SEM. ns, not significant by unpaired two-tailed *t* test. WT *n* = 17, KO *n* = 17 coverslips from 4 independent preparations of neuronal cultures. (*C*) sypHy response at plateau following perfusion with $NH_4Cl$. Scatter plots indicate mean ± SEM. ns, not significant by unpaired two-tailed *t* test. WT *n* = 17, KO *n* = 17 coverslips from 4 independent preparations of neuronal cultures. (*D*) sypHy response from neurons stimulated with 300 APs at 10 Hz (red bar) normalised to the stimulation peak. Neurons were perfused with acidic solution both pre- and post-stimulation for 50 s and 40 s, respectively (yellow bars). (*E*) Rate decay of sypHy fluorescence measured after applying a post-stimulus acidic pulse. Scatter plots indicate mean ± SEM. ns, not significant by unpaired two-tailed *t* test. WT *n* = 16, KO *n* = 13 coverslips from 3 independent preparations of neuronal cultures.

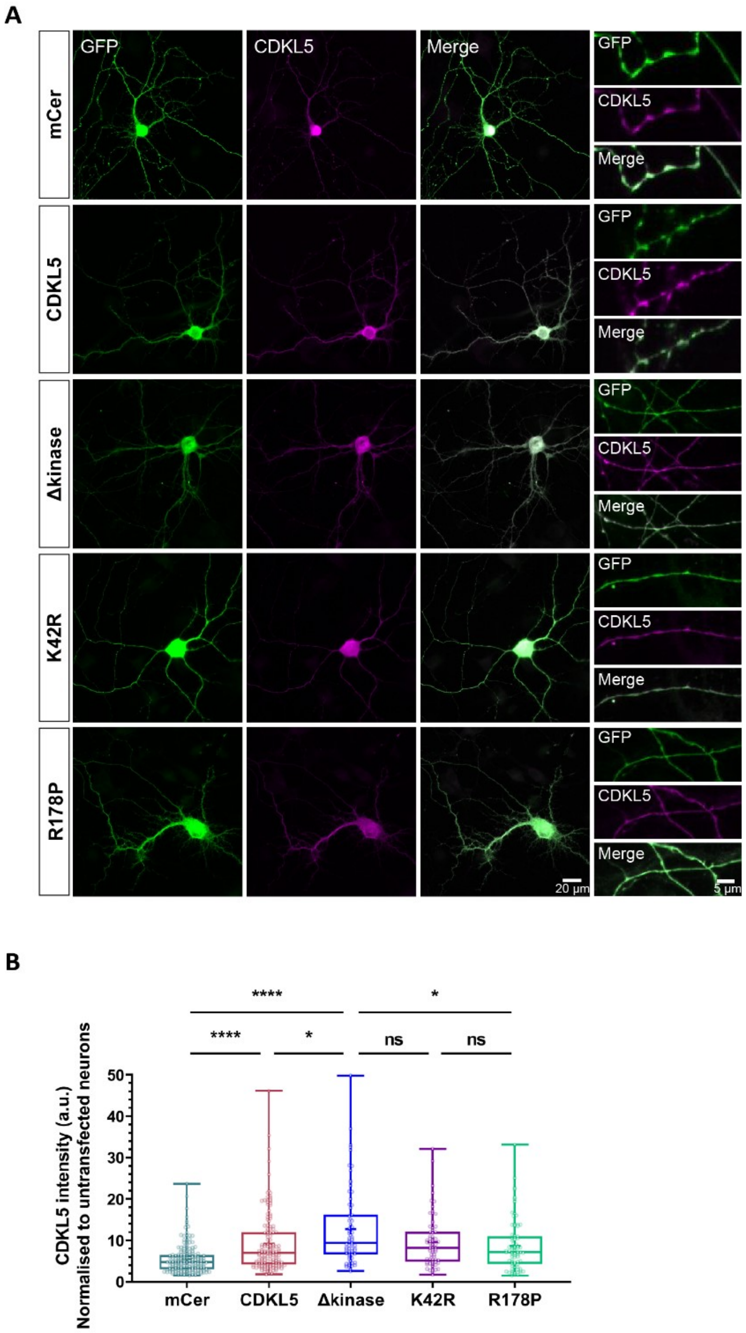


**Figure S3.** GFP-tagged CDKL5 constructs are expressed in primary hippocampal neurons. (*A*) Primary hippocampal neurons from WT rats were transfected with either mCer or mCer-tagged versions of CDKL5 at 8-10 DIV and were fixed at 15 DIV. Representative images of neurons and axons expressing mCer (control) and different CDKL5 constructs labelled for GFP (green) and CDKL5 (magenta). Merged images of GFP and CDKL5. Scale bar, 20 µm (neurons) and 5 µm (axons). (*B*) Quantification of CDKL5 fluorescence intensity of GFP-expressing cell bodies normalised to the intensity of untransfected cell bodies. Box plots present median with IQR indicating min to max whiskers. ns, not significant, $*p < 0.05$, $****p < 0.0001$ by Kruskal-Wallis test followed by Dunn's multiple comparison test, + indicates mean value. mCer $n$ = 191, CDKL5 $n$ = 149, Δkinase $n$ = 71, K42R $n$ = 71, R178P $n$ = 78 cell bodies from 4 independent preparations of neuronal cultures.

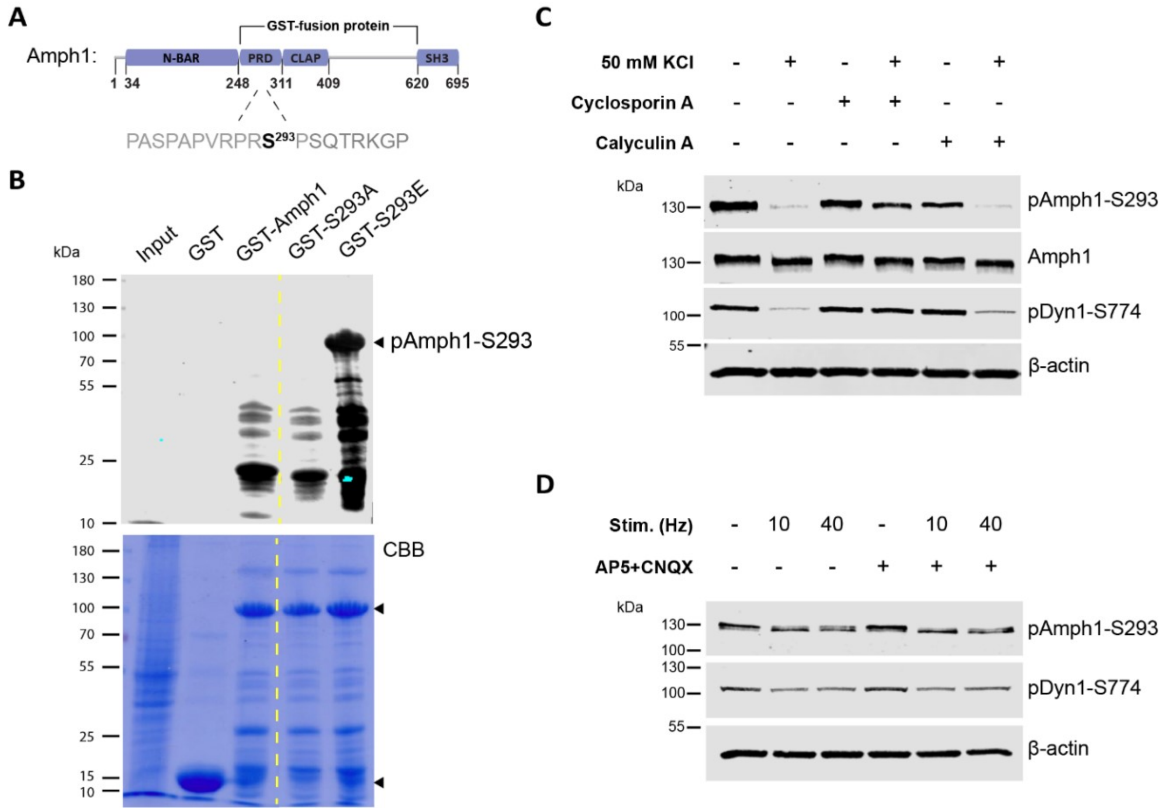


**Figure S4.** Characterisation of a phospho-antibody to report Amph1-S293 phosphorylation dynamics. (*A*) Schematic representation of the structural domains of human Amph1. The residue S293 is located within the PRD of Amph1. (*B*) The GST-fused phosphomutants of Amph1, S293A and S293E, lacking the N-BAR and SH3 domains, as shown in *A*, were generated and expressed in *Escherichia coli*. The pAmph1-S293 antibody detects robustly only the phosphomimetic S293E fusion protein in synaptosomal lysates. All GST-fused proteins were adequately expressed, as the CBB staining indicates (top arrow). Dashed lines indicate cropped images. (*C*) Hippocampal neurons at 14-15 DIV were stimulated with 50 mM KCl for 2 min to trigger neuronal depolarisation resulting in dephosphorylation of Amph1-S293. Treatment with 10 µM cyclosporin A blocks Amph1-S293 dephosphorylation, but treatment with 100 nM calyculin A fails to prevent dephosphorylation at Amph1-S293. The phosphorylation status at Dyn1-S774 was also tested as this site undergoes activity- and calcineurin-dependent dephosphorylation. (*D*) Hippocampal neurons at 14-15 DIV were stimulated with either 300 APs at 10 Hz or 400 APs at 40 Hz in the presence or absence of 50 µM AP5 and 10 µM CNQX. Dephosphorylation at Amph1-S293 occurs at both stimulation frequencies independently of any postsynaptic activity similarly to Dyn1-S774.

| Figure 1 | Group | Mean ± SEM | n = # of fields of view/ N = # of neuronal preparations | Comparison | *p* | Statistical test |
|---|---|---|---|---|---|---|
| Fig. 1C | mCer | 40.42 ± 0.88 | 48/4 | mCer vs. Syp1-mCer | <0.0001 | One-way ANOVA with Tukey's multiple comparison test |
| | Syp1-mCer | 72.99 ± 1.55 | 37/4 | mCer vs. mCer-CDKL5 | 0.0640 | |
| | mCer-CDKL5 | 36.52 ± 1.22 | 32/4 | Syp1-mCer vs. mCer-CDKL5 | <0.0001 | |

| Figure 2 | Group | Mean ± SEM (Fig. 2A) or median (min - max) (Fig. 2C,D) | n = # of neuronal lysates (Fig. 2A) or neurons (Fig. 2C,D)/ N = # of neuronal preparations | Comparison | *p* | Statistical test |
|---|---|---|---|---|---|---|
| Fig. 2A | CDKL5 WT | 1.00 ± 0.10 | 4/4 | WT vs. KO | <0.0001 | Unpaired two-tailed t test |
| | CDKL5 KO | 0.02 ± 0.01 | 4/4 | | | |
| | CHC WT | 1.00 ± 0.09 | 4/4 | | 0.4921 | |
| | CHC KO | 0.85 ± 0.19 | 4/4 | | | |
| | Dyn1 WT | 1.00 ± 0.12 | 4/4 | | 0.1808 | |
| | Dyn1 KO | 0.79 ± 0.06 | 4/4 | | | |
| | Syndapin 1 WT | 1.00 ± 0.25 | 4/4 | | 0.6711 | |
| | Syndapin 1 KO | 0.87 ± 0.13 | 4/4 | | | |
| | Endophilin A1 WT | 1.00 ± 0.28 | 4/4 | | 0.4505 | |
| | Endophilin A1 KO | 0.74 ± 0.16 | 4/4 | | | |
| | VGLUT1 WT | 1.00 ± 0.07 | 4/4 | | 0.9899 | |
| | VGLUT1 KO | 0.10 ± 0.21 | 4/4 | | | |
| | ATP6V1B2 WT | 1.00 ± 0.12 | 4/4 | | 0.2114 | |
| | ATP6V1B2 KO | 0.83 ± 0.04 | 4/4 | | | |
| | Syp1 WT | 1.00 ± 0.07 | 4/4 | | 0.0592 | |
| | Syp1 KO | 1.33 ± 0.12 | 4/4 | | | |
| | pAkt-S473 WT | 1.00 ± 0.07 | 4/4 | | 0.6186 | |
| | pAkt-S473 KO | 1.05 ± 0.05 | 4/4 | | | |
| | pGSK3α/β-S9/S21 WT | 1.00 ± 0.08 | 4/4 | | 0.2271 | |
| | pGSK3α/β-S9/S21 KO | 1.21 ± 0.13 | 4/4 | | | |
| Fig. 2C | $SV2A^+$ WT | 19.67 (4.33 - 43.33) | 144/4 | WT vs. KO | 0.2854 | Mann Whitney two-tailed test |
| | $SV2A^+$ KO | 19.00 (4.33 - 45.00) | 142/4 | | | |
| Fig. 2D | $VGLUT1^+$ WT | 22.83 (8.00 - 45.67) | 144/4 | WT vs. KO | 0.2302 | Mann Whitney two-tailed test |
| | $VGLUT1^+$ KO | 21.50 (9.67 - 41.67) | 142/4 | | | |

| Figure 3 | Group | Mean ± SEM | n = # of coverslips/ N = # of neuronal preparations | Comparison | *p* | Statistical test |
|---|---|---|---|---|---|---|
| Fig. 3C | WT | 0.44 ± 0.02 | 12/4 | WT vs. KO | 0.8932 | Unpaired two-tailed t test |
| | KO | 0.45 ± 0.02 | 12/4 | | | |
| Fig. 3D | WT | 0.16 ± 0.04 | 12/4 | WT vs. KO | 0.0022 | Unpaired two-tailed t test |
| | KO | 0.36 ± 0.04 | 12/4 | | | |
| Fig. 3F | WT | 0.49 ± 0.02 | 13/4 | WT vs. KO | 0.3025 | |

| | | | | | | | |
|---|---|---|---|---|---|---|---|
| | KO | | 0.53 ± 0.03 | 14/4 | | | Unpaired two-tailed t test |
| Fig. 3G | WT | | 0.22 ± 0.03 | 13/4 | WT vs. KO | 0.0065 | Unpaired two-tailed t test |
| | KO | | 0.32 ± 0.02 | 14/4 | | | |
| Fig. 3I | WT | | 0.45 ± 0.03 | 12/4 | WT vs. KO | 0.6015 | Unpaired two-tailed t test |
| | KO | | 0.48 ± 0.05 | 13/4 | | | |
| Fig. 3J | WT | | 0.15 ± 0.03 | 12/4 | WT vs. KO | 0.0442 | Unpaired two-tailed t test |
| | KO | | 0.24 ± 0.02 | 13/4 | | | |

| **Figure 4** | **Group** | | **Mean ± SEM** | **n = # of coverslips/ N = # of neuronal preparations** | **Comparison** | ***p*** | **Statistical test** |
|---|---|---|---|---|---|---|---|
| Fig. 4B | WT | Empty | 0.04 ± 0.03 | 15/4 | WT Empty vs. KO Empty<br>WT Empty vs. WT-CDKL5<br>WT Empty vs. KO-CDKL5<br>KO Empty vs. WT-CDKL5<br>KO Empty vs. KO-CDKL5<br>WT-CDKL5 vs. KO-CDKL5 | 0.0419<br>0.9644<br>0.9956<br>0.0134<br>0.0229<br>0.9947 | Two-way ANOVA with Tukey's multiple comparison test |
| | | CDKL5 | 0.01 ± 0.02 | 14/4 | | | |
| | KO | Empty | 0.17 ± 0.04 | 16/4 | | | |
| | | CDKL5 | 0.03 ± 0.04 | 15/4 | | | |
| Fig. 4D | WT | Empty | 0.10 ± 0.02 | 20/5 | WT Empty vs. KO Empty<br>WT Empty vs. WT-CDKL5<br>WT Empty vs. KO-CDKL5<br>KO Empty vs. WT-CDKL5<br>KO Empty vs. KO-CDKL5<br>WT-CDKL5 vs. KO-CDKL5 | 0.0158<br>0.9982<br>0.5756<br>0.0317<br>0.3186<br>0.7013 | Two-way ANOVA with Tukey's multiple comparison test |
| | | CDKL5 | 0.10 ± 0.02 | 18/5 | | | |
| | KO | Empty | 0.20 ± 0.03 | 21/5 | | | |
| | | CDKL5 | 0.14 ± 0.02 | 19/5 | | | |

| **Figure 5** | **Group** | **Mean ± SEM** | **n = # of coverslips/ N = # of neuronal preparations** | **Comparison** | ***p*** | **Statistical test** |
|---|---|---|---|---|---|---|
| Fig. 5C | KO | 0.26 ± 0.01 | 17/4 | KO vs. CDKL5<br>KO vs. K42R<br>KO vs. R178P | 0.0006<br>0.5993<br>0.6845 | One-way ANOVA with Dunnett's multiple comparison test |
| | CDKL5 | 0.14 ± 0.02 | 16/4 | | | |
| | K42R | 0.23 ± 0.03 | 15/4 | | | |
| | R178P | 0.23 ± 0.02 | 15/4 | | | |
| Fig. 5E | KO | 0.34 ± 0.03 | 13/4 | KO vs. CDKL5<br>KO vs. K42R<br>KO vs. R178P | 0.0135<br>0.9827<br>0.8653 | One-way ANOVA with Dunnett's multiple comparison test |
| | CDKL5 | 0.22 ± 0.02 | 14/4 | | | |
| | K42R | 0.33 ± 0.03 | 13/4 | | | |
| | R178P | 0.31 ± 0.03 | 14/4 | | | |

| **Figure 6** | **Group** | **Mean ± SEM** | **n = # of coverslips/ N = # of neuronal preparations** | **Comparison** | ***p*** | **Statistical test** |
|---|---|---|---|---|---|---|
| Fig. 6C | KO | 0.19 ± 0.02 | 13/4 | KO vs. CDKL5<br>KO vs. ΔC<br>KO vs. Δkinase | 0.0044<br>0.0010<br>0.4857 | One-way ANOVA with Dunnett's multiple comparison test |
| | CDKL5 | 0.07 ± 0.03 | 11/4 | | | |
| | ΔC | 0.07 ± 0.02 | 18/4 | | | |
| | Δkinase | 0.15 ± 0.02 | 15/4 | | | |
| Fig. 6E | KO | 0.25 ± 0.02 | 11/3 | KO vs. CDKL5<br>KO vs. ΔC<br>KO vs. Δkinase | 0.0331<br>0.0281<br>0.9334 | One-way ANOVA with Dunnett's multiple comparison test |
| | CDKL5 | 0.17 ± 0.02 | 10/3 | | | |
| | ΔC | 0.16 ± 0.02 | 10/3 | | | |
| | Δkinase | 0.26 ± 0.02 | 10/3 | | | |

| Figure 7 | Group | | Mean ± SEM | n = # of coverslips/ N = # of neuronal preparations | Comparison | *p* | Statistical test |
|---|---|---|---|---|---|---|---|
| Fig. 7C | WT | Rest | 1.00 ± 0.09 | 3/3 | WT vs. KO overall<br><br>WT vs. KO Rest<br>WT vs. KO KCl<br>WT vs. KO Repol. 2.5 min<br>WT vs. KO Repol. 5 min<br>WT vs. KO Repol. 10 min | 0.5341<br><br>0.9332<br>>0.9999<br>0.9864<br>0.9990<br>0.5856 | Two-way ANOVA with Sidak's multiple comparison test |
| | | KCl | 0.03 ± 0.02 | 3/3 | | | |
| | | Repol. 2.5 min | 0.80 ± 0.23 | 3/3 | | | |
| | | Repol. 5 min | 0.81 ± 0.04 | 3/3 | | | |
| | | Repol. 10 min | 0.76 ± 0.21 | 3/3 | | | |
| | KO | Rest | 0.85 ± 0.05 | 3/3 | | | |
| | | KCl | 0.06 ± 0.03 | 3/3 | | | |
| | | Repol. 2.5 min | 0.69 ± 0.06 | 3/3 | | | |
| | | Repol. 5 min | 0.75 ± 0.06 | 3/3 | | | |
| | | Repol. 10 min | 1.03 ± 0.22 | 3/3 | | | |
| Fig. 7F | WT | DMSO-Rest | 1.00 ± 0.16 | 3/3 | WT vs. KO overall<br><br>WT vs. KO DMSO-Rest<br>WT vs. KO DMSO-KCl<br>WT vs. KO DMSO-Repol.<br>WT vs. KO Inhibitors-Rest<br>WT vs. KO Inhibitors-KCl<br>WT vs. KO Inhibitors-Repol. | 0.8115<br><br>0.9977<br>0.9999<br>0.9967<br>0.9094<br>>0.9999<br>0.9624 | Two-way ANOVA with Sidak's multiple comparison test |
| | | DMSO-KCl | 0.17 ± 0.02 | 3/3 | | | |
| | | DMSO-Repol. | 0.79 ± 0.03 | 3/3 | | | |
| | | Inhibitors-Rest | 0.69 ± 0.06 | 3/3 | | | |
| | | Inhibitors-KCl | 0.05 ± 0.03 | 3/3 | | | |
| | | Inhibitors-Repol. | 0.12 ± 0.05 | 3/3 | | | |
| | KO | DMSO-Rest | 0.94 ± 0.13 | 3/3 | | | |
| | | DMSO-KCl | 0.13 ± 0.06 | 3/3 | | | |
| | | DMSO-Repol. | 0.85 ± 0.13 | 3/3 | | | |
| | | Inhibitors-Rest | 0.81 ± 0.10 | 3/3 | | | |
| | | Inhibitors-KCl | 0.05 ± 0.03 | 3/3 | | | |
| | | Inhibitors-Repol. | 0.02 ± 0.01 | 3/3 | | | |

| Figure S2 | Group | Mean ± SEM | n = # of coverslips/ N = # of neuronal preparations | Comparison | *p* | Statistical test |
|---|---|---|---|---|---|---|
| Fig. S2B | WT | 2.41 ± 0.14 | 17/4 | WT vs. KO | 0.3494 | Unpaired two-tailed t test |
| | KO | 2.63 ± 0.18 | 17/4 | | | |
| Fig. S2C | WT | 0.39 ± 0.01 | 17/4 | WT vs. KO | 0.3477 | Unpaired two-tailed t test |
| | KO | 0.41 ± 0.01 | 17/4 | | | |
| Fig. S2E | WT | 0.52 ± 0.04 | 16/3 | WT vs. KO | 0.5061 | Unpaired two-tailed t test |
| | KO | 0.48 ± 0.03 | 13/3 | | | |

| Figure S3 | Group | Median (min - max) | n = # of cell bodies/ N = # of neuronal preparations | Comparison | *p* | Statistical test |
|---|---|---|---|---|---|---|
| Fig. S3B | mCer | 4.74 (1.53 - 23.67) | 191/4 | mCer vs. CDKL5<br>mCer vs. Δkinase<br>mCer vs. K42R<br>mCer vs. R178P<br>CDKL5 vs. Δkinase<br>CDKL5 vs. K42R<br>CDKL5 vs. R178P<br>Δkinase vs. K42R<br>Δkinase vs. R178P<br>K42R vs. R178P | <0.0001<br><0.0001<br><0.0001<br><0.0001<br>0.0119<br>>0.9999<br>>0.9999<br>0.9257<br>0.0272<br>>0.9999 | Kruskal-Wallis test with Dunn's multiple comparison test |
| | CDKL5 | 6.96 (1.81 - 46.13) | 149/4 | | | |
| | Δkinase | 9.34 (2.59 - 49.79) | 71/4 | | | |
| | K42R | 8.25 (1.75 - 32.12) | 71/4 | | | |
| | R178P | 7.15 (1.49 - 33.12) | 78/4 | | | |

**Table S1.** Table of experimental *n*, *p* values and statistical tests.
